# A cryptic transactivation domain of EZH2 binds AR and AR’s splice variant promoting oncogene activation and tumorous transformation

**DOI:** 10.1101/2022.08.04.502794

**Authors:** Jun Wang, Kwang-Su Park, Xufen Yu, Weida Gong, H. Shelton Earp, Gang Greg Wang, Jian Jin, Ling Cai

**Affiliations:** Lineberger Comprehensive Cancer Center, University of North Carolina at Chapel Hill School of Medicine, Chapel Hill, NC 27599, USA; Department of Biochemistry and Biophysics, University of North Carolina at Chapel Hill School of Medicine, Chapel Hill, NC 27599, USA; Mount Sinai Center for Therapeutics Discovery, Departments of Pharmacological Sciences and Oncological Sciences, Tisch Cancer Institute, Icahn School of Medicine at Mount Sinai, New York, NY 10029, USA; Department of Pharmacology, University of North Carolina at Chapel Hill School of Medicine, Chapel Hill, NC 27599, USA; Department of Medicine, University of North Carolina at Chapel Hill School of Medicine, Chapel Hill, NC, 27599, USA; Department of Genetics, University of North Carolina at Chapel Hill School of Medicine, Chapel Hill, NC 27599, USA

**Keywords:** EZH2, PRC2, AR, AR-V7, histone modification, prostate cancer, inhibitor, PROTAC

## Abstract

Enhancer of Zeste Homolog 2 (EZH2) and androgen receptor (AR) are crucial chromatin regulators involved in the development and/or progression of prostate tumor, including advanced castration-resistant prostate cancer (CRPC). To sustain prostate tumorigenicity, EZH2 establishes noncanonical biochemical interaction with AR for mediating oncogene activation, in addition to its canonical role as a transcriptional repressor and enzymatic subunit of Polycomb Repressive Complex 2 (PRC2). However, the molecular basis underlying non-canonical activities of EZH2 in prostate cancer remains elusive and therapeutic strategies for targeting EZH2:AR-mediated oncogene activation activities are also lacking. Here, we report that a cryptic transactivation domain of EZH2 (EZH2^TAD^) binds both AR and AR spliced variant 7 (AR-V7, an AR variant enriched in CRPC), mediating assembly and/or recruitment of transactivation-related machineries at genomic sites that lack PRC2 binding. Such noncanonical targets of EZH2:AR/AR-V7:(co)activators are enriched for the clinically-relevant oncogenes. We also show that EZH2^TAD^ is required for the chromatin recruitment of EZH2, for EZH2-mediated oncogene activation, and for CRPC growth in vitro and in vivo. To completely block EZH2’s multifaceted oncogenic activities in prostate cancer, we employed MS177, a recently developed proteolysis targeting chimera (PROTAC) of EZH2. Strikingly, MS177 achieved on-target depletion of both EZH2’s canonical (EZH2:PRC2) and noncanonical (EZH2^TAD^:AR/AR-V7:coactivators) complexes in prostate tumor, eliciting much more potent antitumor effects than the catalytic inhibitors of EZH2. Overall, this study reports previously unappreciated requirements of EZH2^TAD^ for mediating EZH2’s noncanonical (co)activator recruitment and gene-activation functions in prostate tumor and suggests EZH2-targeting PROTACs as potentially attractive therapeutics for the treatment of aggressive prostate tumors that rely on the circuits wired by EZH2 and AR.

## Introduction

Prostate cancer is the most frequently diagnosed, non-cutaneous malignancy in men, causing approximately 30,000 deaths annually in USA(1). Androgen deprivation therapy (ADT) is the standard treatment of prostate cancer(2); yet it is almost inevitable that patients receiving ADT become refractory and eventually develop castration-resistant prostate cancer (CRPC)(3,4). Both genetic and epigenetic deregulations play critical roles during the development of prostate tumor and its stepwise progression into more advanced, more aggressive forms, notably CRPC. Androgen receptor (AR) and Enhancer of Zeste Homolog 2 (EZH2) are among the most relevant prostate tumor-promoting oncoproteins. Amplification of the AR gene, mutation of AR’s ligand-binding domain (LBD) and/or cofactor-directed mechanisms can lead to a hyperactive AR signaling during prostate tumorigenesis(5,6); in addition, the AR gene undergoes alternative splicing with inclusion of cryptic exons in CRPC, giving rise to the truncated, hormone-independent, constitutively active AR variants such as AR splice variant 7 (AR-V7)(7,8).

EZH2 is widely known as the catalytic subunit of Polycomb Repressive Complex 2 (PRC2), which induces trimethylation of histone H3 lysine 27 (H3K27me3) to maintain repression of a battery of tumor-suppressive, differentiation-promoting, and/or immunity-related transcripts(9–14). Overexpression of EZH2 due to genomic loss of EZH2-targeting microRNA is associated with prostate tumor progression, correlating to poor clinical outcomes(15). Increasing evidence demonstrated that, in prostate cancers including CRPC, EZH2 also has nonconventional functions in binding to non-PRC2 partners(16) such as AR and fibrillarin, for potentiating transcriptional activation(17–20) and translation(21), respectively. Studies of other tumors such as breast cancer also showed EZH2 over-expression not correlated with the increase of H3K27me3(22), consistent to the PRC2-independent role of EZH2. Indeed, recent study of MLL-rearranged acute myeloid leukemia (AML)(23) demonstrated that EZH2 directly binds cMyc and p300 through a cryptic transactivation domain (TAD), independently of its PRC2 function, to promote oncogenesis(23,24). However, the role for the transactivation domain of EZH2 (EZH2^TAD^) in prostate tumor remains unexplored to date.

Previously, it has been shown that knockdown (KD) or knockout (KO) of EZH2 in various prostate tumor models suppressed cancer cell proliferation(15, 17–19,21), laying a strong foundation for therapeutically targeting EZH2 as an attractive strategy for the treatment of prostate cancer. Small-molecule inhibitors that selectively target the methyltransferase activity harbored within EZH2’s Su(var)3-9, Enhancer-of-zeste and Trithorax (SET) domain have been developed and are currently under clinical development(15,25). However, it is most likely that these EZH2 enzymatic inhibitors cannot block EZH2’s methyltransferase-independent functions such as those related to gene activation and/or scaffolding (e.g. recruiting/binding non-PRC2 factors), some of which have been suggested to be equally critical for oncogenesis(18,21,23). Strategies for completely blocking the multilevel activities of EZH2 in prostate tumor need to be developed.

## Materials and Methods

### Cell Lines

Cell lines used in the study included 293T (American Tissue Culture Collection [ATCC], CRL-3216), 22Rv1 (ATCC, CRL-2505), LNCaP (ATCC, CRL-1740), C4-2 (ATCC, CRL-3314), PC3 (ATCC, CRL-1435), DU145 (ATCC, HTB-81) and RWPE-1 (ATCC, CRL-11609), which were cultured and maintained according to the vendor-provided protocol. Cell lines of LNCaP-abl, LNCaP_RB-/-/P53-/- were kindly provided by Dr. Zoran Culig (Innsbruck Medical University, Austria) and Dr. Charles Sawyers (MSKCC) respectively. Identity of cell lines was ensured by UNC’s Tissue Culture Facility with genetic signature analyses and examination of mycoplasma contamination performed using commercial detection kits (Lonza, #LT27-286).

### Chemicals

MS177, C24, MS177N1 and MS177N2 were synthesized and used as described previously(23). Pomalidomide (#36471) was purchased from AstaTech. GSK126 (#S7061), EPZ6438 (#S7128), MLN4924 (#S7109) and A-485 (#S8740) were purchased from Selleck Chemical. CPI-1205 and UNC1999 were synthesized as previously reported(39,40). UNC6852 (EED PROTAC) was used as before(23).

### Antibodies

Antibodies used in the work included mouse anti-AR-V7 (Precision, Cat # AG10008), mouse anti-EZH2 (BD, Cat # 612666), rabbit anti-H3K27me3 (Millipore, Cat # 07-449), Sheep anti-EED (R&D, Cat # AF5827), mouse anti-M2 Flag tag (sigma, Cat #F1804), GST-HRP (GeneTex, Cat # GTX114099), anti-full-length (FL) AR (against AR-FL C-terminus, C19) (Santa Cruz, Cat # sc-815), rabbit anti-AR (against AR N-terminus, N20) (Santa Cruz, Cat # sc-816), mouse anti-Ubiquitin (Santa Cruz, Cat # SC-8017), rabbit anti-H3K27ac (Abcam, Cat # ab4729), rabbit anti-H3 (Abcam, Cat # ab1791), rabbit anti-EZH2 (Cell Signaling Technology, Cat # 5246), rabbit anti-H3K27me3 (Cell Signaling Technology, Cat # 9733), rabbit anti-HA tag (Cell Signaling Technology, Cat # 3724), rabbit anti-GAPDH (Cell Signaling Technology, Cat # 5174), rabbit anti-SUZ12 (Abcam, Cat # ab12073), rabbit anti-AR (Cell Signaling Technology, Cat # 5153), beta-Actin (Cell Signaling Technology, Cat # 4970), rabbit anti-CRBN (against AR N-terminus, Cell Signaling Technology, Cat # 71810), rabbit anti-PARP (Cell Signaling Technology, Cat # 9532), rabbit anti-Cleaved Caspase-3 (Cell Signaling Technology, Cat # 9661), rabbit anti-Cleaved Caspase-7 (Cell Signaling Technology, Cat # 8438), and normal rabbit IgG (Cell Signaling Technology, Cat # 2729). HRP-linked secondary antibodies, either anti-mouse IgG (Cat # 7076) or anti-rabbit IgG (Cat # 7074), were obtained from Cell Signaling Technology.

### Cleavage Under Targets & Release Using Nuclease (CUT&RUN)

CUT&RUN was performed as previously described(23). Briefly, 0.5 million of cells were first collected, washed in the CUT&RUN wash buffer, and then bound to the activated ConA beads (Bangs Laboratories, #BP531). Next, the cell:bead sample was incubated with antibodies against the protein target (1:100 dilution) and then permeabilized in the digitonin-containing buffer, which was then followed by washing in the digitonin buffer, incubation with pAG-MNase, and another washing in the digitonin buffer to remove the unbound pAG-MNase. After the final wash, cells were subjected to digestion following the pAG-MNase activation by addition of the pAG-MNase digestion buffer, followed by incubation on rotator for 2 hours at 4 °C. Solubilized chromatin was then released using the CUT&RUN stop buffer, in which equal amount of Drosophila spike-in chromatin (0.5 ng/sample) was added for spike-in normalization, and DNA purification was carried out with the PCR cleanup kit. About 5 ng of the purified CUT&RUN DNA was used for preparation of multiplexed libraries with the NEB Ultra II DNA Library Prep Kit per manufacturer’s instruction. Sequencing was conducted using an Illumina NextSeq 500 Sequencing System.

### Chromatin Immunoprecipitation (ChIP) Followed by PCR (ChIP-qPCR)

ChIP-qPCR were performed as previously described(23,27,49). Primers used for ChIP-qPCR are listed in Supplementary Table S4.

### ChIP Sequencing (ChIP-seq) and CUT&RUN Data Analysis

ChIP-seq data downloaded from NCBI Gene Expression Omnibus (GEO) were re-analyzed as previously described(23). For CUT&RUN, raw reads were mapped to the reference genome (hg19) using bowtie v2.3.5. The non-primary alignment, PCR duplicates, or blacklist regions were removed from aligned data by Samtools (v1.9), Picard ‘MarkDuplicates’ function (v2.20.4), and bedtools (v2.28.0), respectively. Peak calling was performed using MACS2 (macs2 callpeak -f BAMPE -g hs/mm -keep-dup 1). Deeptools (v3.3.0) was used to generate bigwig files. Genomic binding profiles were generated using the deepTools ‘bam-Compare’ functions. Profiles of CUT&RUN read densities were displayed in Integrative Genomics Viewer (IGV, Broad Institute). Heatmaps for ChIP-seq signals were generated using the deepTools “computeMatrix” and “plotHeatmap” functions. Distribution of peaks were analyzed by ‘annotatePeaks.pl’ function of HOMER (Hypergeometric Optimization of Motif Enrichment)(50). Motif analysis was performed by the SeqPos tool in Cistrome to uncover the motifs that are enriched close to the peak centers by taking the peak locations as the input(51).

### RNA Sequencing (RNA-seq) and Data Analysis

RNA-seq was performed as described(23,27). Total RNA was first purified using RNeasy Plus Mini Kit (Qiagen, #74136) and then treated with Turbo DNA-free kit (Thermo, #AM1907) to remove genomic DNA. Multiplexed RNA-seq libraries were subjected for deep sequencing. Reads were mapped to the reference genome followed by analysis of differentially expressed genes (DEG) as before(27,52). Fastq files were aligned to the GRCh38 human genome (GRCh38.d1.vd1.fa) using STAR v2.4.2(53) with parameters: --outSAMtype BAM Unsorted --quantMode TranscriptomeSAM. Transcript abundance was estimated with salmon v0.1.19(54) to quantify the transcriptome defined by Gencode v22. Gene level counts were summed across isoforms and genes with low counts (maximum expression < 10) were filtered for the downstream analyses. Raw read counts were used for differential gene expression analysis by DESeq2 v1.38.2(55) where size normalization factor was estimated based on median-of-ratios. GSEA(56) was performed as described(23,52). Expression heatmaps were generated using mean-centered log2 converted TPM (Transcripts Per Million) sorted in descending order based on expression values in R’s package “gplots” v3.0.3 with either no clustering or column hierarchical clustering by average linkage. Volcano plots visualizing DEGs were produced using R’s package “EnhancedVolcano” v3.11. Annotation of DEGs was conducted using Metascape(57). DisGeNET(58) was used to annotate genes associated with human disease.

### Plasmid Construction

Wildtype (WT) or the serially deleted version of AR was fused in-frame with Flag tag and subsequently cloned into the mammalian expression vector of pcDNA3.1 (Invitrogen). GST fusion constructs with full-length EZH2 or EZH2^TAD^, either WT or TAD-transactivation-dead mutant (including F171A+F145A [FA] or F171K+145K [FK]), and the pCDH-EF1-neo lentiviral constructs containing Flag-tagged EZH2 (WT or TAD-transactivation-dead mutant) were previously described and used as before(23). EZH2’s SET domain deletion was generated by a site-directed mutagenesis kit (Agilent, 200521). All plasmids were verified by Sanger sequencing. Primers for making constructs were listed in Supplementary Table S4.

### Transient Transfection

293T cells were seeded in 100 mm dish with fresh medium, and then transfected on the next day with PEI (sigma) following manufacturer’s instructions. Cells were harvested 48 hours after transfection for analysis such as co-IP or immunoblotting.

### The siRNA- or shRNA-mediated Gene Knockdown (KD)

Cells were transfected with siRNA using the Lipofectamine RNAi MAX reagent according to manufacturer’s instruction. The MISSION® esiRNA for human CRBN (sigma, # EHU047571) was ordered and used per vendor’s guideline. The pLKO.1-based lentiviral shRNA plasmids for KD of EZH2 (23), AR or ARV7 (27) were described before.

### Viral Production and Stable Cell Line Generation

Lentiviruses were prepared with the packaging system in 293T cells. In brief, 293T cells were co-transfected with lentiviral vector and the packaging plasmids (psPAX2 and pMD 2.5G) and the supernatant containing viruses were harvested at 48- and 72-hours post-transfection. After filtration with 0.45-μM filters, viruses were used to infect target cells in the presence of 8 μg/mL polybrene. 48 hours post-infection, cells were selected with either 1 μg/mL of puromycin (Gibco) or 1 mg/mL of Geneticin (Gibco) for 7 days to establish stable expression cell lines.

### Immunoblotting (IB)

Cells were collected and lysed in EBC buffer (50 mM Tris pH 8.0, 120 mM NaCl, 0.5% NP40, 0.1 mM EDTA, and 10% glycerol) freshly supplemented with a complete protease inhibitor cocktail (Roche) and phosphatase inhibitor (Roche). Protein concentration of cell lysates was measured by Bradford assay (BioRad). Equal amounts of protein lysates were separated by SDS-PAGE and transferred to PVDF membranes (Millipore). Quantification of the intensity of protein bands by normalizing to GAPDH was performed by ImageJ software.

### Co-immunoprecipitation (co-IP) and GST Pulldown

IP was performed as described previously(23,59). Cell pellets were lysed in EBC buffer (freshly supplemented with protease/phosphatase inhibitors) on ice for 30 minutes. After sonication, debris was removed by centrifugation at 12,000 g for 15 minutes at 4 °C. For IP of tagged protein, HA-conjugated (Roche, #11815016001) beads were incubated with lysate overnight at 4 °C. GST tagged protein were purified as previously described(23). GST pulldown was conducted with cell lysate and 1 μg of GST-fusion recombinant protein as described(23,59).

### Reverse Transcription Followed by Quantitative Polymerase Chain Reaction (RT-qPCR)

RNA was isolated using the RNeasy PLUS Mini Kit (Qiagen). RT was performed with 1 ug of total RNA using cDNA Reverse Transcription kit according to the manufacturer’s protocols (Invitrogen), followed by qPCR using SYBR Green Master Mix (BioRad) on a QuantStudio 6 Flex Real-Time PCR System (Thermo). The relative abundance of gene expression was calculated using the comparative CT method which compares the Ct value of target gene to that of GAPDH. Primers used for RT-qPCR are listed in Supplementary Table S4.

### Cell Proliferation Assays

Cells were seeded at a density of 1,000-2,500 cells per well in 96-well plates in triplicate and incubated with indicated compounds at different time-points. Fresh medium with compound was changed every two days. At each time-point, MTT reagent (Promega) was added to the cell culture medium and incubated for 1-2 h before being subjected to measure absorbance at 490 nm using CYTATION-5 imaging reader (BioTek). EC_50_ (effective control to 50% growth inhibition) values were calculated using a nonlinear regression analysis of the mean ± SD from at least triplicated datasets for each biological assay.

### 2D Colony Formation Assay

2D-colony formation assay was performed as previously described. Briefly, 5,000 cells were plated on 6-well plates and incubated with medium containing the compound. After 2-3 weeks’ incubation, cells were fixed by 100% methanol and stained with crystal violet.

### Soft Agar Assay

Soft agar-based colony formation assay was performed as described previously(59). Briefly, cells were plated at a density of 24,000 cells/mL for 22Rv1 cells in complete medium supplemented with 0.4% of agarose onto the bottom layers composed of medium with 1% of agarose. Every four days, 0.5 mL of fresh complete media containing compound was added onto the plate. After 3-4 weeks’ incubation, cell culture plates were stained with 100 μg/mL of iodonitrotetrazoliuim chloride solution (Sigma), and after incubation overnight, numbers of cell colonies were counted.

### Cell Fractionation

1 million cells were harvested, washed with cold PBS, and resuspended in 200 μL of CSK buffer (10 mM Pipes pH 7.0, 300 mM sucrose, 100 mM NaCl, 3 mM MgCl_2_, 0.1% Triton X-100, freshly supplemented with protease/phosphatase inhibitor cocktail), followed by incubation on ice for 30 minutes, as described(52). Then, the sample was subject to centrifugation at 1,300 g for 5 minutes at 4 °C to collect supernatant (which contains soluble proteins) and pellet fractions (which contains the chromatin-associated proteins). Cell pellets were dissolved in 1.5x SDS loading buffer. Same amounts of protein sample were used for immunoblotting.

### Ubiquitination Assay

22Rv1 cells treated with different compounds were harvested and extracted in 100 μL of EBC buffer containing 1% of SDS. Cell extract was heat-denatured at 95 °C for 5 min and then diluted with 900 μL of EBC buffer. After brief sonication and centrifugation, lysates were subjected to IP with antibodies of target proteins, followed by anti-ubiquitin immunoblotting.

### Tumor Growth in Xenografted Animal Models

All animal experiments were approved and performed in accord with the guidelines of the Institutional Animal Care and Use Committee (IACUC) at UNC. One million of 22Rv1 cells were suspended in 100 μl with 1:1 mixture of PBS and Matrigel (BD Biosciences) and subcutaneously (s.c.) injected into dorsal flanks of NOD/SCID/gamma-null (NSG) mice bilaterally (carried out by the Animal Studies Core, UNC Lineberger Comprehensive Cancer Center). Diameter measurements of xenografted tumors were performed three times per week using caliper and the tumor volume were calculated. Mice were all sacrificed when the tumors in control group reached the maximum.

### Statistics and Reproducibility

Statistical analyses were performed using GraphPad Prism (version 9). Unpaired two-tailed Student’s t-test was used for experiments comparing two sets of data with assumed normal distribution. Data are presented as mean ± SD from at least three independent experiments. *, **, and *** denote the *P* value of < 0.05, 0.01 and 0.005, respectively. *P* < 0.05 was considered to be statistically significant. NS denotes not significant. No statistical methods were used to predetermine sample size. All data from representative experiments (such as imaging and micrographs) were repeated at least two times independently with similar results.

## Results

### Besides its canonical H3K27me3-cobound EZH2:PRC2 sites, EZH2 also binds genomic sites that are characterized by the gene-activation-associated histone marks and colocalization with RNA polymerase II (Pol II) and AR or AR-V7 in prostate cancer

We sought to systematically define genome-wide binding patterns of EZH2 in prostate cancer by performing <u>C</u>leavage <u>U</u>nder <u>T</u>argets & <u>R</u>elease <u>U</u>sing <u>N</u>uclease (CUT&RUN(26)) for EZH2 and H3K27me3, a gene-repressive histone mark characteristic of canonical EZH2:PRC2 complex, in 22Rv1 cells, a commonly used model of CRPC. Replicated mapping profiles of EZH2 or H3K27me3 were highly correlated (Supplementary Figure S1A), which were additionally correlated to <u>A</u>ssay for <u>T</u>ransposase-<u>A</u>ccessible <u>C</u>hromatin using <u>seq</u>uencing (ATAC-seq) dataset of same cells. A portion of the called EZH2 peaks (8,436 or 34.5%) overlapped H3K27me3 and lacked chromatin accessibility (Figure 1A-B; Supplementary Figure S1B); thus, we termed these EZH2^+^/H3K27me3^+^/low-accessibility sites as EZH2-ensemble or canonical targets of EZH2:PRC2. Meanwhile, a larger proportion of EZH2-binding sites (15,987 or 65.5%) lacked H3K27me3 and, instead, were enriched for high ATAC-seq signals (Figure 1A-B; Supplementary Figure S1B), indicating a feature of open chromatin and a noncanonical function of EZH2; thus, we defined these EZH2^+^/H3K27me3^-^/high-accessibility sites as EZH2-solo targets. Next, we interrogated additional chromatin marks at EZH2-solo sites. Reminiscent of what was observed in MLL-rearranged AML(23), the EZH2-solo peaks identified from 22Rv1 CRPC cells were overwhelmingly enriched for a set of gene-activation-related markers including H3K27ac, H3K4me2, H3K4me3, BRD4 (a histone acetylation reader) and RNA Pol II (Figure 1C, top). Such a correlational pattern was in stark contrast to what was seen with the EZH2^+^/H3K27me3^+^-cobound peaks (Figure 1c, bottom). Furthermore, overall expression of the genes associated with EZH2-solo sites was significantly higher than that with EZH2:PRC2-ensemble peaks, based on RNA-sequencing (RNA-seq) data of 22Rv1 cells(27) (Supplementary Figure S1C).

**Figure 1.**
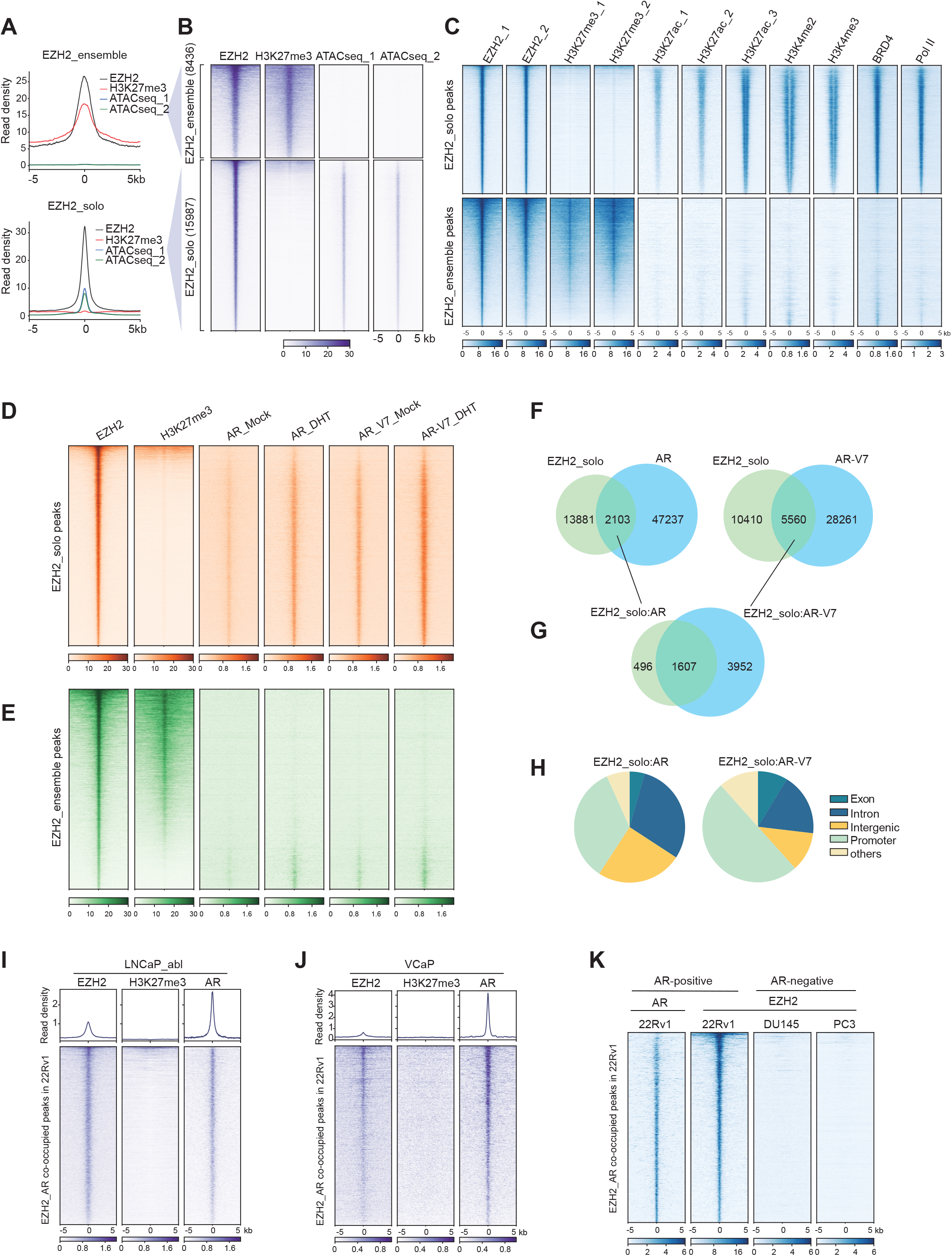
Besides its canonical H3K27me3-cobound EZH2:PRC2 sites, EZH2 also binds genomic sites that are characterized by the gene-activation-associated chromatin markers and colocalization with AR and/or AR-V7 in prostate cancer. **(A-B)** Averaged intensities (**A**) and heatmaps (**B**) for EZH2 or H3K27me3 CUT&RUN and ATAC-seq signals ± 5 kb from the centers of canonical EZH2^+^/H3K27me3^+^/no-accessibility peaks (i.e. EZH2-ensemble, 8,436 peaks; top panels) or noncanonical EZH2^+^/H3K27me3^-^/high-accessibility peaks (i.e. EZH2-solo, 15,987 peaks; bottom panels) in 22Rv1 cells. **(C)** Heatmap for signals of EZH2 (in duplicate), H3K27me3 (in duplicate), H3K27ac (in triplicate), H3K4me2, H3K4me3, BRD4 and RNA Pol II ± 5 kb from the centers of EZH2-solo (top panel) or EZH2:PRC2-ensemble peaks (bottom panel). Except that H3K4me2, H3K4me3 and Pol II data were from LNCaP-abl cells, all the others were generated in 22Rv1 cells. EZH2 and H3K27me3 datasets were from CUT&RUN experiments, and all the others from ChIP-seq. **(D-E)** Heatmap of EZH2, H3K27me3, AR and AR-V7 signals ± 5 kb from the centers of either EZH2-solo (**D**) or EZH2:PRC2-ensemble (**E**) peaks identified in 22Rv1 cells. Mock, ligand-stripped condition; DHT, ligand-treated condition. **(F)** Venn diagram showing the overlapping of EZH2-solo peaks with either AR (left) or AR-V7 (right) in 22Rv1 cells. **(G)** Venn diagram showing a significant overlap between EZH2-solo:AR and EZH2-solo:AR-V7 co-occupied sites in 22Rv1 cells. **(H)** Pie chart showing genomic annotation of EZH2-solo:AR (left) or EZH2-solo:AR-V7 (right) peaks in 22Rv1 cells. **(I-J)** Averaged intensities (top panel) and heatmaps (bottom panel) of EZH2, H3K27me3 and AR ChIP-seq signals, ± 5 kb from the centers of those EZH2-solo:AR co-occupied sites identified from 22Rv1 cells, in LNCaP-abl (**I**) or VCaP (**J**) cells. **(K)** Heatmap of signals of AR (only 22Rv1 cells shown; left) or EZH2 binding, ± 5 kb from the centers of those EZH2-solo:AR co-bound peaks identified in 22Rv1 cells (2^nd^ column), in the AR-negative DU145 and PC3 cells (two right columns).

The above observations highlighted the existence of both classic EZH2:PRC2 sites and noncanonical EZH2-solo targets in prostate cancer, in agreement with previous reports that EZH2 forms interactions with AR, mediating target gene activation instead of repression(17,18,20,28). Consistently, a significant portion of EZH2-solo sites overlapped binding of AR and/or AR-V7 (Figure 1D), both expressed in 22Rv1 cells(27), whereas AR/AR-V7 binding at those canonical PRC2 sites was found to be generally minimal (Figure 1E). Approximately 2,100 of AR sites and 5,500 of AR-V7 sites were EZH2-cobound and absent of H3K27me3, thus termed as EZH2-solo:AR/AR-V7 sites (Figure 1F). In agreement that AR-V7 acts as a constitutively active AR, the majority of EZH2-solo:AR peaks overlapped EZH2-solo:AR-V7 peaks (Figure 1G). Androgen-responsive element (ARE) and the binding motifs of FOXA1, the known cofactor of AR, were most enriched at both EZH2-solo:AR and EZH2-solo:AR-V7 sites (Supplementary Figure S1D-E), indicating an AR/cofactor (FOXA1)-driven recruitment. Gene ontology (GO) and DisGeNET analyses of the sites commonly bound by EZH2-solo, AR and AR-V7 uncovered the enrichment for genes involved in cell proliferation, tissue development, stress response, and prostate cancer metastasis (Supplementary Figure S1F-G). Approximately 50% of EZH2-solo:AR-V7 and 33% of EZH2-solo:AR sites were localized at gene promoters (Figure 1H). To assess whether or not the aforementioned EZH2-solo:AR cobinding is conserved across different prostate cancer models, we next related those EZH2-solo:AR sites identified from 22Rv1 cells with publicly available datasets of various prostate cell lines(18,29). While the EZH2-solo:AR cobound peaks showed similar co-occupancy of both oncoproteins in two additional AR-positive prostate cancer cells, namely LNCaP-abl (Figure 1I) and VCaP cells (Figure 1J), such binding of EZH2 to same genomic sites was lacking in the two AR-negative cells, PC3 and DU145 (Figure 1K). These observations suggested potential recruitment of EZH2 by AR to a subset of AR-binding sites, in a PRC2-independent fashion.

### EZH2, AR and AR-V7 cooperate to activate transcription of a set of the clinically relevant oncogenes in prostate cancer

The above genome-wide profiling unveiled that EZH2-solo sites in prostate cancer exhibited co-localization with AR and/or AR-V7, RNA Pol II and prominent gene-active markers, which pointed to a gene-activation-related role of EZH2, differing from its canonical EZH2:PRC2 function seen at sites exhibiting high H3K27me3 and low chromatin accessibility. To further determine EZH2-mediated gene regulation in prostate cancer, we performed RNA-seq after its KD in 22Rv1 cells (Supplementary Table S1 and Supplementary Figure S2A). As expected, depletion of EZH2 greatly inhibited 22Rv1 cell growth (Supplementary Figure S2B). Analysis of differentially expressed genes (DEGs) showed more transcripts down-regulated than up-regulated upon EZH2 KD relative to control (Supplementary Figure S2C), consistent to more EZH2-binding sites showing chromatin accessibility in 22Rv1 cells (Fig 1A-C). GO analysis of DEGs down-regulated following EZH2 depletion showed the enrichment of pathways related to cell cycle, DNA replication, and cell metabolism (Supplementary Figure S2D). Furthermore, we compared the transcriptomic profiles of 22Rv1 cells after EZH2 KD with those following either AR-specific or AR-V7-specific KD(27). Here, we identified 130 genes to be co-activated by AR-FL, AR-V7 and EZH2 (Figure 2A, Supplementary Figure S2E, Supplemental Table S2). These EZH2:AR:AR-V7 co-upregulated genes were generally bound directly by EZH2, AR, AR-V7, coactivators (BRD4), RNA Pol II and gene-active histone marks, but absent of H3K27me3 (Figure 2B), which again pointed to gene-activation effects by EZH2 and AR/AR-V7 at these targets. GO (Figure 2C) and DisGeNET (Figure 2D) analyses also showed the EZH2-solo:AR:AR-V7 co-activated genes enriched for the pathways associated with cell cycle, DNA repair, metabolism, and prostate cancer progression and metastasis. Through integration of genomic binding and RNA-seq profiles, we determined genes that were co-upregulated by EZH2, AR and AR-V7 and also exhibited direct co-bindings of the three (Figure 2E, labeled on the right), as exemplified by *PRC1, CDK2, MYBL2, HMGB1* and *PDIA4* (Figure 2F and Supplementary Figure S2F). High expression of EZH2-solo:AR:AR-V7 co-activated transcripts were found correlated with poor prognosis of prostate cancer- the patients with higher expression of *PRC1, CDK2* or *MYBL2* in tumors exhibited the significantly worse clinical outcomes (Figure 2G-H); additionally, the expression of these genes was positively associated with the EZH2 and AR levels in the TCGA prostate cancer patients (Supplementary Figure S2G). Reverse transcription followed by quantitative PCR (RT-qPCR) confirmed that the high expression of select EZH2-solo:AR:AR-V7 target genes indeed relied on the presence of AR, AR-V7 and EZH2 (Supplementary Figure S2H-J), but was generally unaffected (or even slightly increased) upon treatment with C24, a potent EZH2 enzymatic inhibitor (which completely suppressed H3K27me3), or UNC6852, a PROTAC degrader of EED(30) (which depleted the EED-associated PRC2 and thus H3K27me3) (Figure 2I, purple and green; Supplementary Figure S2K-L). In stark contrast, expression of the same genes was significantly inhibited by treatment with the p300/CBP catalytic inhibitor A-485, which suppressed histone acetylation(31) (Figure 2i; Supplementary Figure S2M). Furthermore, compared to mock, KD of AR or AR-V7 significantly reduced the direct EZH2 binding to EZH2-solo:AR:AR-V7 target sites (Figure 2J); as a control, AR or AR-V7 loss did not affect EZH2 protein levels in these cells (Supplementary Figure S2N). Thus, establishment of EZH2-solo binding relies on the presence of AR/AR-V7, in agreement with a lack of EZH2-solo:AR:AR-V7 sites in the AR-negative prostate tumor cells (Figure 1K). Together, our results substantiated a PRC2-independent function of EZH2 at its solo targets in prostate cancer, which acts in concert with AR and/or AR-V7 for target gene activation, with potential involvement of p300/CBP and BRD4 as coactivators.

**Figure 2.**
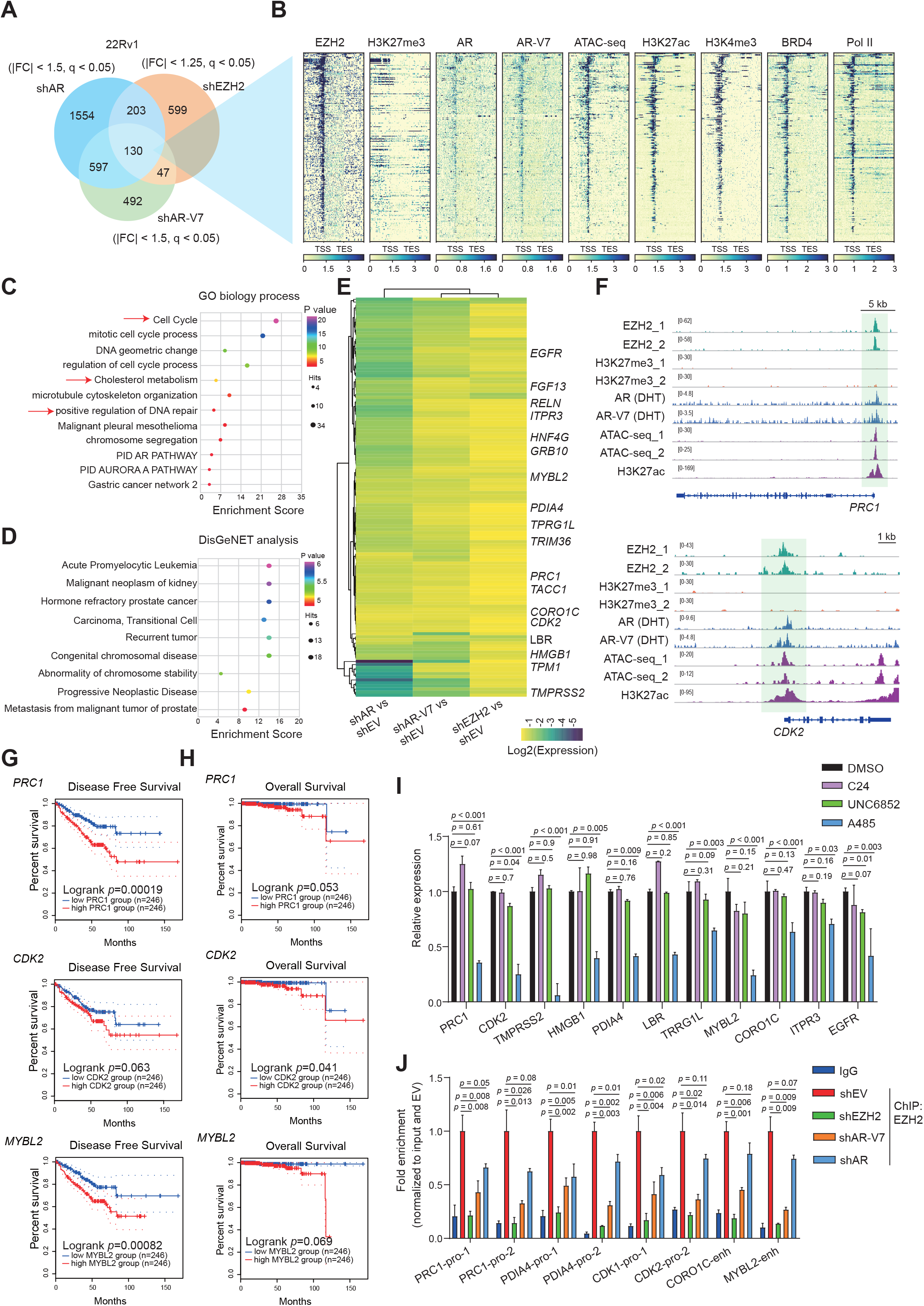
EZH2, AR and AR-V7 cooperate to activate transcription of a set of the clinically relevant oncogenes in prostate cancer. **(A)** Venn diagram using DEGs, identified by RNA-seq to be downregulated in 22Rv1 cells post-depletion of either EZH2, AR or AR-V7 (n = 2 biologically independent experiments). FC, fold change; q, adjusted *P* value. **(B)** Heatmap showing overall binding of the indicated protein at the EZH2:AR:AR-V7 co-upregulated genes (n = 130; defined in **a**). TSS, transcriptional start site; TES, transcriptional end site. **(C-D)** Gene Ontology (GO) analysis (**c**) and enrichment of the DisGeNet category (**d**) using the EZH2:AR:AR-V7 co-upregulated genes in **a**. **(E)** Heatmap using the indicated RNA-seq sample comparison shows log2-converted ratios of the 130 genes co-upregulated by EZH2, AR and AR-V7 in 22Rv1 cells. Genes with direct co-bindings by EZH2, AR and AR-V7 are labelled at the right side. EV, empty vector. **(F)** Integrative Genomics Viewer (IGV) image of enrichment for the indicated factor at the *PRC1* (top) or *CDK2* gene (bottom) in 22Rv1 cells. **(G-H)** Kaplan-Meier disease-free survival (**G**) and overall survival (**H**) analysis based on the *PRC1* (top), *CDK2* (middle) or *MYBL2* (bottom) expression in patient samples from the prostate cancer cohort of The Caner Genomics Atlas (TCGA). Statistical significance was determined by log-rank test and shown. **(I)** RT-qPCR for the indicated EZH2:AR:AR-V7 co-upregulated genes in 22Rv1 cells, treated with 2.5 μM of UNC6852, C24 or A-485, relative to DMSO, for 24 hours. The y-axis shows averaged signals after normalization to those of GAPDH and to mock-treated (n = 3; mean ± s.d.; unpaired two-tailed Student’s t-test). **(J)** ChIP-qPCR for EZH2 binding at the promoter (pro) or enhancer (enh) of indicated EZH2:AR:AR-V7 co-targeted genes in 22Rv1 cells after depletion of EZH2, AR or AR-V7, relative to mock (red). The y-axis shows averaged signals after normalization to those of input and then to control samples (n = 3; mean ± s.d.; unpaired two-tailed Student’s t-test). IgG (dark blue; using shEV samples) serves as a non-specific antibody control.

### Interaction between EZH2^TAD^ and AR is required for establishment of EZH2-solo binding at AR sites and for the malignant growth of prostate cancer

EZH2 associates with AR(17,18,20,32), and we validated interactions of EZH2 with AR and AR variants using both co-immunoprecipitation (co-IP; Figure 3A, lane 2 vs 1) and glutathione S-transferase (GST) pulldown (Supplementary Figure S3A). The biochemical basis underlying such EZH2:AR/variant interactions, however, remains murky. Recently, a cryptic transactivation domain of EZH2 (EZH2^TAD^) was reported to interact directly with cMyc and coactivators (such as p300) for promoting target gene activation(23,24,28). We thus queried whether or not EZH2^TAD^ also interacts with AR/variants in prostate cancer. First, GST pulldown using recombinant EZH2^TAD^ readily detected interaction with both AR and AR variant, expressed either endogenously or exogenously (Figure 3B and Supplementary Figure S3B; lane 2 vs 1), and such AR/variant interactions to EZH2^TAD^ were completely abolished by the EZH2^TAD^ mutation that carries substitution of the two key hydrophobic residues (Phe145 and Phe171) with either alanine or charged lysine residue, which dramatically reduced the transactivation effect by EZH2^TAD^ (23,24) (Figure 3B and Supplementary Figure S3B; see lanes of FA or FK vs WT). In agreement, full-length AR interacted with both wildtype (WT) and SET-domain-deleted forms of EZH2, but not the two EZH2^TAD^-transactivation-dead mutants, in co-IP (Figure 3A; lanes 3 and 4 vs. 2 and 5), supporting an essential requirement of EZH2^TAD^ for mediating EZH2:AR interaction.

**Figure 3.**
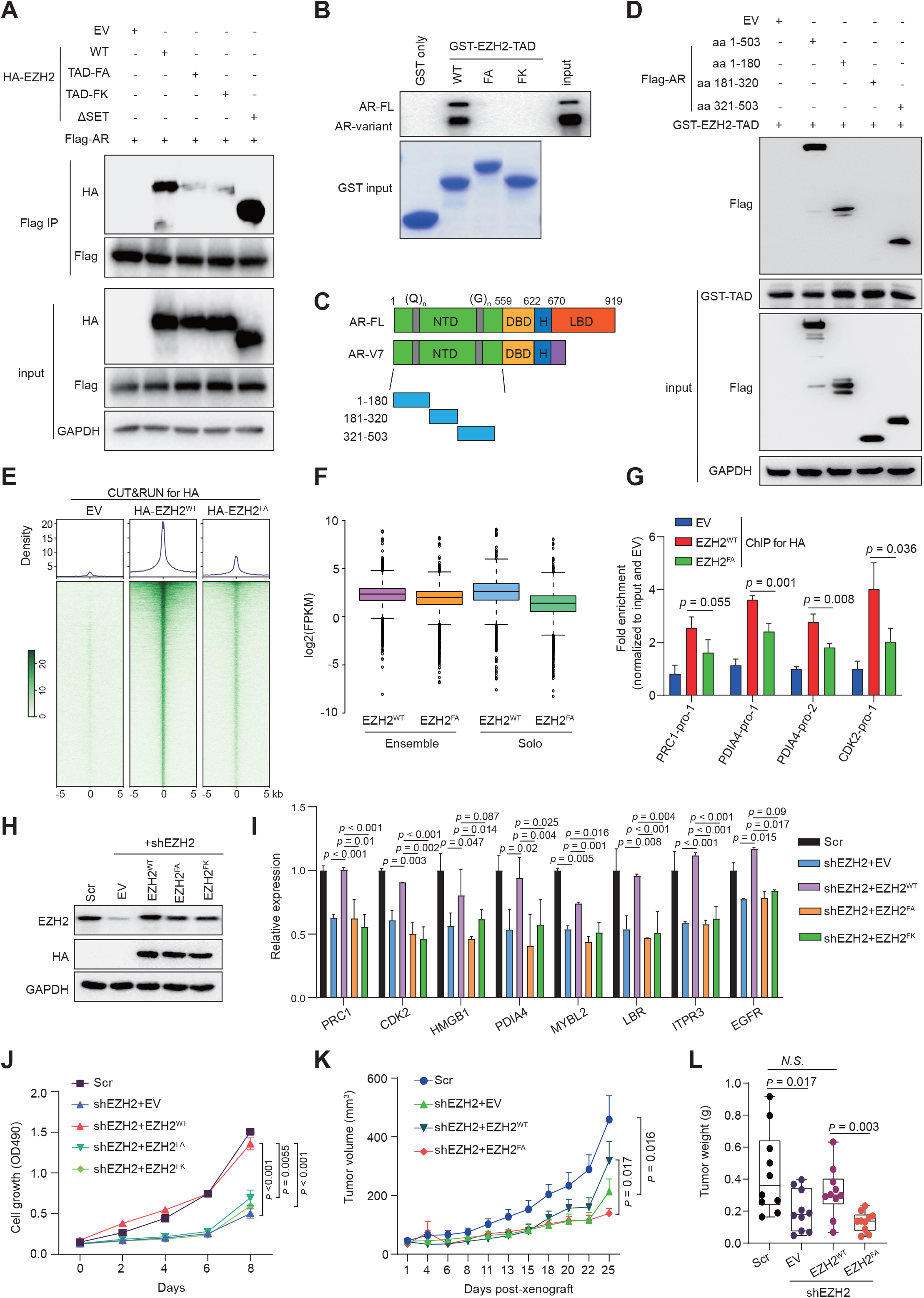
Interaction between EZH2^TAD^ and AR is required for establishment of EZH2-solo binding at AR sites and for the malignant growth of prostate cancer cells. **(A)** Anti-Flag immunoprecipitation (IP; top panel) for Flag-tagged AR and the indicated HA-tagged EZH2, either wildtype (WT) or with the indicated EZH2^TAD^ mutation (FA or FK) or the SET deletion, in 293T cells. FA, F145A+F171A; FK, F145K+F171K. **(B)** GST pulldown using the indicated GST protein, either GST alone or fused with WT or mutant EZH2^TAD^, and the total lysate of 22Rv1 cells, followed by anti-AR immunoblotting (top). AR-FL, full-length AR. **(C)** Scheme of the domain organization of AR-FL and AR-V7. NTD, N-terminal domain; DBD, DNA-binding domain; LBD, ligand-binding domain; (Q)_n_, poly-Q region; (G)_n_, poly-G region. **(D)** Pulldown using GST-tagged WT EZH2^TAD^ and total lysate of 293T cells transfected with the indicated Flag-tagged AR, followed by anti-Flag immunoblotting (top). **(E)** Averaged intensities (top) and heatmap (bottom) for EZH2 binding (exogenously expressed HA-tagged EZH2, either WT or TAD-mutated [FA], as assayed by anti-HA CUT&RUN), ± 5 kb from the center of the called peaks in 22Rv1 cells. Cells with empty vector (EV) served as CUT&RUN control. **(F)** Boxplot showing log2-transformed FPKM counts of EZH2, WT or TAD-mutated, at EZH2-solo (right) or EZH2:PRC2-ensemble (left) sites in 22Rv1 cells. The boundaries of box plots indicate the 25th and 75th percentiles, the center line indicates the median, and the whiskers (dashed) indicate 1.5× the interquartile range. **(G)** ChIP-qPCR of binding by HA-EZH2, either WT or TAD-mutant (FA), at the indicated EZH2-solo:AR co-targeted genes. The y axis shows signals after normalization to those of input (n = 3; mean ± s.d.; unpaired two-tailed Student’s t-test). EV-transduced cells served as a negative control. **(H-J)** Immunoblotting for the level of rescued EZH2 (**H**), RT-qPCR (**I**) for EZH2-solo:AR:AR-V7 coactivated genes, and measurement for 22Rv1 cell growth (**J**) after mock treatment (scramble; Scr) or depletion of endogenous EZH2 (shEZH2) in 22Rv1 cells, pre-rescued with exogenous shEZH2-resistant HA-EZH2, either WT or TAD-mutant (FA or FK). For **i** and **j**, n = 3, mean ± s.d.; unpaired two-tailed Student’s t-test. **(K-L)** Averaged volume (**K**) and weight (**L**) in xenografted tumors by using 22Rv1 cells, which were pre-rescued with exogenous shEZH2-resistant HA-EZH2, either WT or TAD-mutant, and then subjected for mock treatment (scramble or Scr) or endogenous EZH2 KD. Tumor weight was measured at day 25 post-xenograft in NSG mice (n = 10, mean ± s.d.; unpaired two-tailed Student’s t-test).

We next queried what region within AR mediates EZH2^TAD^ interaction. Here, we found that GST-EZH2^TAD^ bound both full-length and C-terminal-truncated forms of AR (Supplementary Figure S3C), showing that the AR N-terminal domain (amino acids 1-503) is sufficient in mediating EZH2^TAD^ interaction. Furthermore, we constructed a set of serial deletion constructs for GST pulldown and further narrowed down the EZH2^TAD^-interacting regions to two AR segments—its amino acids 1-180, which is enriched for glutamines (polyQ), and amino acids 321-503, which is enriched for glycines (polyG) (Figure 3C-D).

To discern the role for EZH2^TAD^ in mediating genomic recruitment of EZH2, we conducted CUT&RUN for WT or TAD-mutated EZH2 following its stable expression into 22Rv1 cells. Compared to WT, the TAD-mutated EZH2 exhibited significantly reduced genomic binding (Figure 3E), as exemplified by what was seen at *PRC1* and *MYBL2* (Supplementary Figure S3D). Notably, there was a preferential decrease of binding at EZH2-solo regions compared to that at EZH2:PRC2 ensemble regions (Figure 3F, right vs left). ChIP-qPCR corroborated that, relative to WT, the EZH2^TAD^ mutant exhibited the significantly reduced binding at the tested EZH2-solo targets (Figure 3G). Lastly, we evaluated the requirement of EZH2^TAD^ for prostate cancer growth. Using 22Rv1 cells with endogenous EZH2 depleted (Figure 3H), we found that, compared to WT EZH2, its TAD-transactivation-dead mutants failed to rescue the decreased expression of EZH2 target genes (Figure 3I) and failed to rescue the EZH2-loss-related growth inhibition in vitro (Figure3J and Supplementary Figure S3E). Meanwhile, EZH2’s SET domain was also found to be essential for sustaining 22Rv1 cell proliferation (Supplementary Figure S3F-G), consistent to previous reports supporting an involvement of EZH2:PRC2 during prostate oncogenesis(33,34). Additionally, WT but not the TAD-transactivation-dead EZH2 rescued the defect in the 22Rv1 xenografted tumor growth caused by loss of endogenous EZH2 (Figure 3K-I).

Overall, our observations demonstrated the previously unappreciated, indispensable requirements of EZH2^TAD^ for mediating EZH2 interaction with AR/variants, for EZH2’s chromatin recruitment to EZH2-solo sites, and for promoting prostate tumor growth in vitro and in vivo.

### MS177, the EZH2-targeting PROTAC, mediates on-target degradation of EZH2, as well as EZH2-associated canonical (EZH2:PRC2) and non-canonical (EZH2:AR/AR-V7) complexes, in prostate cancer

A handful of small molecules have been developed to block the catalytic activity harbored within EZH2’s SET domain, some of which are under clinical evaluation(33). However, we predicted efficacies of these EZH2 enzymatic inhibitors to be low due to failure in targeting noncanonical, SET(PRC2)-independent functions of EZH2(23), such as EZH2:AR-mediated prostate oncogene activation, which are TAD-dependent and also clinically relevant. This notion was corroborated by a generally mild or a lack of effect by C24, a potent EZH2 SET inhibitor(35), on growth of prostate cancer cells in vitro (Supplementary Figure S4A). Using Proteolysis Targeting Chimera (PROTAC) technology, we recently developed MS177, an EZH2-targeting degrader, that can simultaneously deplete EZH2 and EZH2-associated factors in cells(23). Thus, we examined the EZH2-degrading effect by MS177 in prostate tumor cells. As expected, MS177 treatment in 22Rv1 cells dose-dependently depleted cellular EZH2, regardless of being soluble or chromatin-associated (Supplementary Figure S4B), and its PRC2 partners (EED and SUZ12), an effect not observed following comparable treatment with C24 or either of the two inactive analogue compounds of MS177, namely MS177N1 (E3 binding-dead) and MS177N2 (C24 binding-dead)(23) (Figure 4A). The half-maximal degradation concentration (DC50) value of MS177 in 22Rv1 cells was measured to be 0.86 ± 0.12 μM, and maximum degradation (D_max_) value 77% (Supplementary Figure S4c), supporting that MS177 is indeed a valuable tool for degrading EZH2 in prostate cancer. More excitingly, we observed that, unlike the PROTAC-inactive controls (C24, MS177N1 or MS177N2), MS177 also concentration-dependently degraded both AR and/or AR-V7 across multiple tested lines of prostate cancer including 22Rv1, C4-2 and LNCaP (Figure 4B-C, and Supplementary Figure S4D). The effects by MS177 on degradation of EZH2:PRC2 and AR/AR-V7 were found generally comparable (Figure 4B-C, and Supplementary Figure S4D). In contrast, the EED degrader UNC6852 efficiently depleted EED and EED-associated EZH2:PRC2 but did not alter the AR/AR-V7 levels (Figure 4D), consistent with a PRC2-independent association of EZH2 with AR/variants as we observed at EZH2-solo sites.

**Figure 4.**
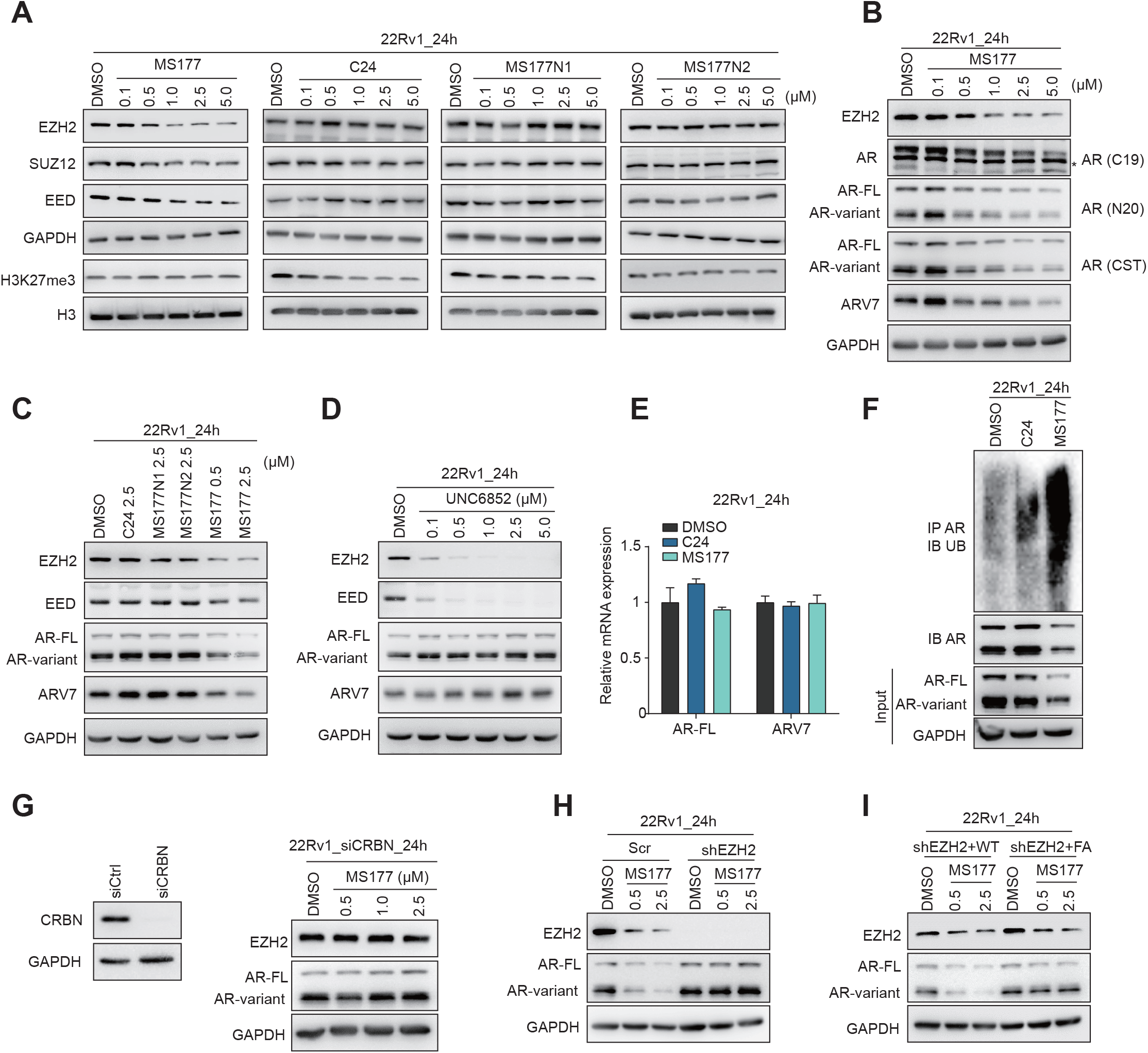
MS177, the EZH2-targeting PROTAC, mediates on-target degradation of EZH2, as well as EZH2-associated canonical (EZH2:PRC2) and non-canonical (EZH2:AR/AR-V7) complexes, in prostate cancer. **(A)** Immunoblotting of PRC2 subunits (EZH2, EED and SUZ12) and H3K27me3 in 22Rv1 cells treated with the indicated concentration of MS177, C24, MS177N1 or MS177N2 versus DMSO, for 24 hours. **(B)** Immunoblotting of EZH2, full-length AR (AR-FL) and AR-V7 (probed with different antibodies recognizing either AR-FL or AR-V7 only or both isoforms) in 22Rv1 cells treated with the indicated concentration of MS177, versus DMSO, for 24 hours. *, nonspecific band. **(C-D)** Immunoblotting of EZH2, EED, AR and AR-V7 in 22Rv1 cells treated with 2.5 μM of C24, MS177N1, MS177N2 or different concentrations of MS177 **(C)**, or different indicated concentrations of an EED-targeting PROTAC, UNC6852 **(D)**, versus DMSO, for 24 hours. **(E-F)** RT-qPCR **(E)** for AR-FL and AR-V7 mRNA levels and immunoblots **(F)** examining AR ubiquitination in 22Rv1 cells post-treatment with DMSO or 2.5 μM of C24 or MS177 for 24 hours. The y-axis in **(E)** shows averaged signals after normalization to GAPDH and to mock-treated (n = 3; mean ± s.d.). Top panels in **(F)** used samples from anti-AR pulldown followed by immunoblotting with anti-ubiquitin antibodies, and bottom panels showed input sample immunoblotting. **(G)** Left: Immunoblotting of CRBN in 22Rv1 cells transfected with either control (siCtl) or CRBN-targeting siRNA (siCRBN). Right: Immunoblots for EZH2 and AR in 22Rv1 cells transfected with CRBN-targeting siRNA, followed by treatment with the indicated concentration of MS177, versus DMSO, for 24 hours. **(H-I)** Immunoblotting of EZH2 and AR in 22Rv1 cells expressing either scramble (Scr) or EZH2-targeting (shEZH2) shRNA **(H)**, or in those EZH2-depleted 22Rv1 cells rescued with exogenously-expressed WT or TAD-mutated (FA) EZH2 **(I)**, followed by treatment with different doses of MS177 for 24 hours compared to DMSO.

We next explored MS177’s mechanism of action (MOA) for AR/variant degradation. First, cellular AR was found ubiquitinated and then depleted post-treatment with MS177 but not C24, while the transcriptional levels of AR/AR-V7 stayed unchanged (Figure 4E-F), suggesting a proteasome-dependent protein degradation mechanism underlying AR depletion. Additionally, the MS177-induced EZH2 degradation was effectively blocked by pre-treatment of cells with pomalidomide (the E3 ligase ligand module of MS177; Supplementary Figure S4E) or MLN4924 (a NEDD8 activation and neddylation inhibitor that suppresses assembly of Cullin-based E3 ligase^32^; Supplementary Figure S4F). Depletion of CRBN, the E3 ligase that MS177 recruits, also almost completely abrogated MS177-induced depletion of both EZH2 and AR/variant (Figure 4G vs 4B). Lastly, depletion of EZH2 completely blocked the MS177-mediated degradation of AR/variant (Figure 4H, right vs. left), and in 22Rv1 cells with endogenous EZH2 depleted, MS177 was able to degrade the TAD-mutated form of EZH2 (harboring the intact SET, to which MS177 binds) but not AR/variant anymore (Figure 4I), supporting that the AR/variant degradation by MS177 requires the presence of EZH2 and the EZH2^TAD^-directed binding of AR.

Together, MS177, an EZH2-targeting PROTAC, effectively degrades EZH2, as well as the EZH2-associated canonical (EZH2:PRC2) and non-canonical (EZH2:AR/AR-V7) complexes in prostate cancer, which has not been reported before.

### Genomics profiling further substantiates the effects of MS177 on inhibiting both EZH2:PRC2- and AR/AR-V7-related oncogenic nodes

To further delineate gene-regulatory effects of MS177 in prostate tumor, we performed spike-in-controlled CUT&RUN for EZH2, H3K27m3 and AR (using the N-20 antibody that recognizes both full-length AR and AR variants) after treatment of 22Rv1 cells with MS177. Compared to mock, MS177 significantly decreased genome-wide binding of EZH2, both at EZH2:PRC2-ensemble sites and at EZH2-solo sites (Figure 5A-B; Supplementary Figure S5A), exemplified by what was seen at noncanonical EZH2-solo targets such as *PRC1, MYBL2* and *CDK2* (Figure 5C and Supplementary Figure S5B) and classic EZH2:PRC2 targets such as *CCND2, MYT1* and *WNT2B* (Figure 5D and Supplementary Figure S5C). As expected, MS177 treatment also decreased overall H3K27me3 level from EZH2:PRC2 sites (Figure 5E-F and Supplementary Figure S5D; for examples, see also bottom panels of Figure 5D and Supplementary Figure S5C). Relative to mock, MS177 treatment caused the expected decrease of AR binding at EZH2-solo sites where AR cobound (Figure 5G-H); more interestingly, MS177’s inhibitory effect on AR binding was found to be genome-wide and extended to those AR sites where EZH2 does not bind at all (Supplementary Figure S5E), as observed at classic targets of AR/variant such as *KLK3, KLK2* and *FKBP5* (Figure 5I and Supplementary Figure S5F). Such a genome-wide decrease of AR binding is likely due to a ‘drainage’ effect by MS177 on global AR proteins in cells (see also Discussion section), highlighting a potential advantage of the EZH2-targeting PROTAC degrader (MS177) over conventional enzymatic inhibitors of EZH2.

**Figure 5.**
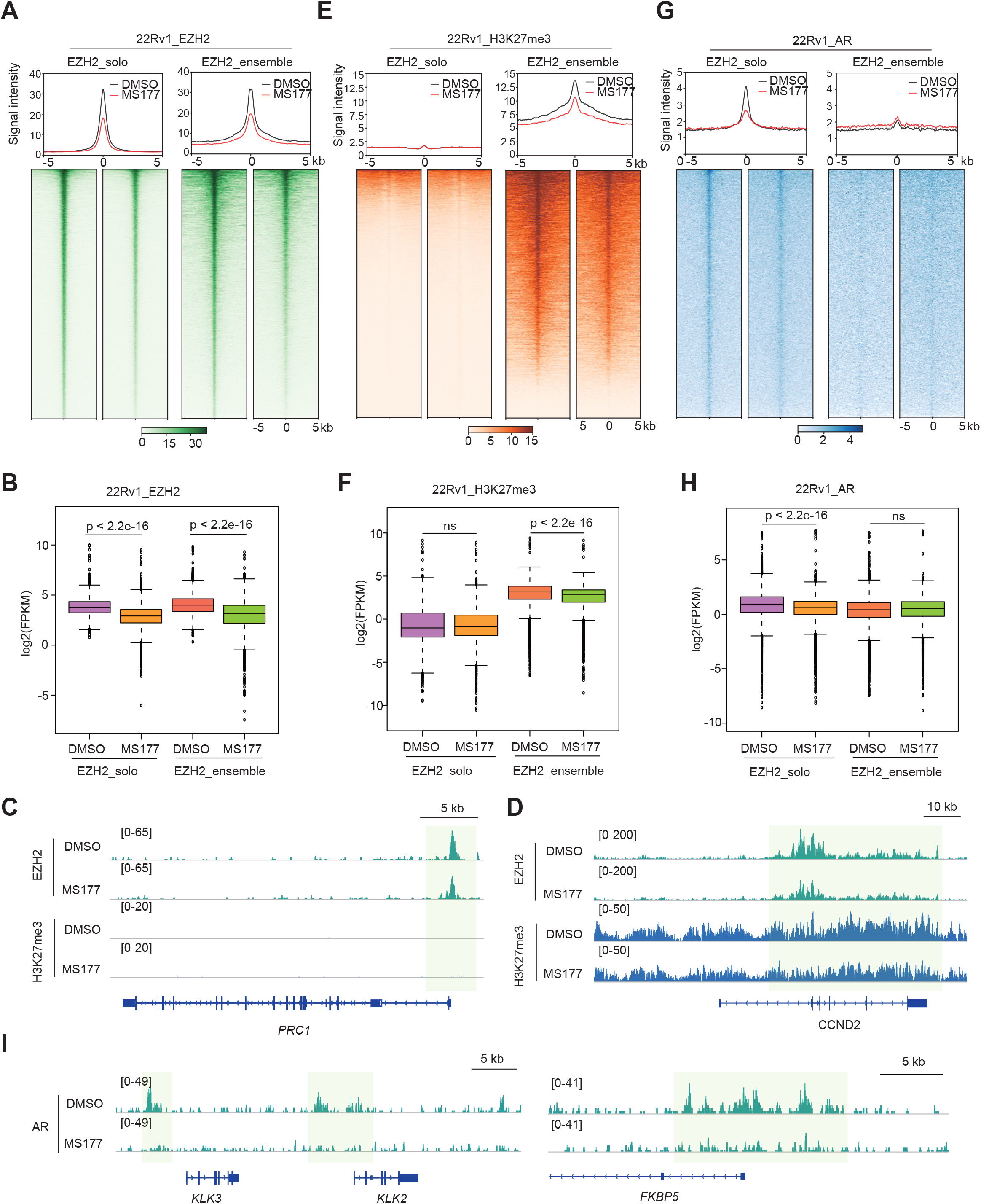
CUT&RUN-based profiling demonstrates effects by MS177 on decreasing genomic binding of both EZH2 and AR/AR-V7 in prostate cancer. **(A,E,G)** Average intensities (top panel) and heatmaps (bottom panel) showing CUT&RUN signals of EZH2 (**A**), H3K27me3 (**E**) and AR (**G**) (normalized against spikein control and sequencing depth) ±□5□kb around the peak centers of EZH2-solo (left) or EZH2:PRC2-ensemble sites (right) identified in 22Rv1 cells, treated for 24 hours with DMSO (black) or 2.5 μM of MS177 (red). **(B,F,H)** Log2-transformed RPKM counts for CUT&RUN signals of EZH2 (**B**), H3K27me3 **(F)** and AR (**H**) at those EZH2-solo (left) or EZH2:PRC2-ensemble sites (right) identified in 22Rv1 cells, treated for 24 hours with DMSO or 2.5 μM of MS177. **(C-D)** IGV views of EZH2 and H3K27me3 binding (spike-in control and depth normalized) at *PRC1* **(C)** or *CCND2* **(D)** post-treatment of 22Rv1 cells with DMSO or 2.5 μM of MS177. **(I)** IGV views of AR binding (spike-in control and depth normalized) at *KLK3 and KLK2* (left) and *FKBP5* (right) post-treatment of 22Rv1 cells with DMSO or 2.5 μM of MS177.

Next, we evaluated transcriptome-modulatory effects by MS177, compared to its inactive control compounds. Here, we treated 22Rv1 cells with either DMSO, MS177, C24 or MS177N1, followed by RNA-seq (Supplementary Table S3). Dramatic transcriptomic changes were observed only after treatment with MS177, but not C24 or MS177N1 (Figure 6A). Those MS177-activated DEGs only showed mild changes following comparable treatment with C24 or MS177N1 (Figure 6B). Likewise, those EZH2-repressed genes, defined as DEGs re-activated upon EZH2 KD, were found de-repressed by treatment with MS177, and not C24 or MS177N1 (Figure 6C), again demonstrating a superior/unique effect of MS177 on altering transcriptome of prostate tumor. Gene Set Enrichment Analysis (GSEA) revealed treatment of MS177 associated with re-activation of known H3K27me3- or PRC2-repressed genes, reminiscent of what was seen with RNA-seq profiles of cells with EZH2 KD (Figure 6D; Supplementary Figure S6A). Meanwhile, the AR/AR-V7 signaling genes were found significantly downregulated upon MS177 treatment relative to mock (Figure 6E-G; Supplementary Figure S6B). Moreover, those genes coactivated by EZH2, AR and AR-V7, as we defined by KD studies (Figure 2A), were significantly inhibited by treatment with MS177, and not C24 or MS177N1 (Figure 6H). By RT-qPCR, we further confirmed the unique effect by MS177 on downregulating the select EZH2-solo:AR/AR-V7 target genes in 22Rv1 cells (Figure 6I).

**Figure 6.**
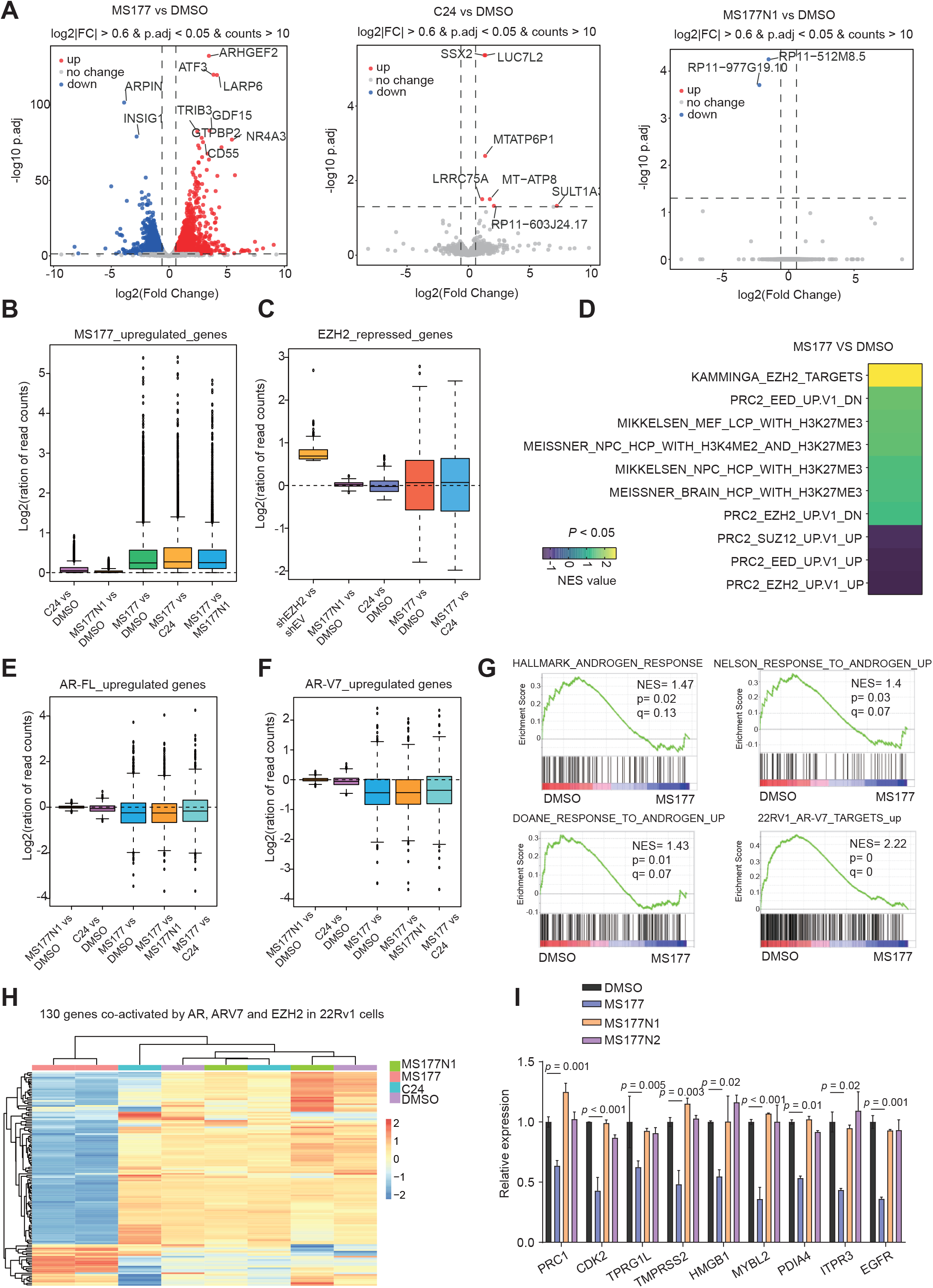
RNA-seq further substantiates unique and superior effects by MS177 on inhibiting both EZH2:PRC2- and AR/AR-V7-related oncogenic programs in prostate cancer. **(A)** Volcano plots showing transcriptomic alterations in 22Rv1 cells following treatment with 2.5 μM of MS177 (left), C24 (middle) or MS177N1 (right), compared to DMSO, for 24 hours. DEGs with significant expression changes were highlighted. **(B-C)** Box plots showing the log2 ratios for DEGs upregulated in 22Rv1 cells treated with MS177 versus DMSO **(B)**, or DEGs upregulated after EZH2 KD in 22Rv1 cells (i.e., EZH2-repressed genes; **C)**. Comparison was indicated on x-axis, including C24 vs DMSO, MS177N1 vs DMSO, MS177 vs DMSO, and MS177 vs C24. **(D)** Heatmap of GSEA normalized enrichment score (NES) values revealing that MS177 treatment is highly correlated with de-repression of the PRC2- or H3K27me3-repressed genes. **(E-F)** Box plots showing the log2 ratios for DEGs significantly downregulated in 22Rv1 cells after AR (**E**) or AR-V7 (**F**) KD relative to control. Comparison was indicated on x-axis, including C24 versus vs DMSO, MS177N1 vs DMSO, MS177 vs DMSO, MS177 vs MS177N1 and MS177 vs C24. **(G)** GSEA revealing that, relative to control, MS177 treatment in 22Rv1 cells is correlated with downregulation of the indicated genesets known to be activated by AR or AR-V7. **(H)** Heatmap showing the expression of 130 genes co-activated by EZH2, AR and AR-V7 in 22Rv1 cells, treated with MS177, C24 or MS177N1, compared to DMSO. Treatment condition was indicated by colored bars on the top of heatmap, with the legend shown on the right. **(I)** RT-qPCR for the indicated EZH2:AR:AR-V7 co-activated target genes in 22Rv1 cells, treated with DMSO or 2.5 μM of C24, MS177N1 or MS177 for 24 hours. Y-axis shows RT-qPCR signals after normalization to those of GAPDH and then to DMSO-treated cells (n = 3 independent experiments; presented as the mean ± SD). For panels **B, C, E** and **F**, the boundaries of box plots indicate the 25th and 75th percentiles, the center line indicates the median, and the whiskers (dashed) indicate 1.5× the interquartile range.

Collectively, our integrated CUT&RUN and RNA-seq profiling following pharmacological treatment or genetic depletion of EZH2 lent strong support for on- target effects by MS177, resulting in simultaneous suppression of both EZH2:PRC2 and AR/AR-V7-directed oncogenic circuits in prostate cancer.

### Compared to the EZH2 enzymatic inhibitors, MS177 elicits much more potent antitumor effects in prostate cancer cells

Next, we sought to evaluate antitumor effects of MS177 in a panel of cell lines representing different stages of prostate cancer, and half-maximal effective concentration (EC_50_) values of MS177 were measured (Figure 7A). First, MS177 had little growth-inhibitory effect in a nonmalignant prostate epithelial line RWPE1 (Figure 7A bottom; Supplementary Figure S7A), suggesting that MS177 is not generally cytotoxic. In contrast, MS177 demonstrated consistent and fast-acting anti-proliferation effects in all examined prostate cancer cell models (Figure 7A-C; Supplementary Figure S7B-C). Notably, MS177 exhibited a magnitude increase of efficacy in inhibiting growth of 22Rv1 and LNCaP cells, compared to its non-PROTAC controls, C24, MS177N1 or MS177N2 (Figure 7B-D; see also Supplementary Figure S4A for C24). We also confirmed the MS177-induced EZH2 degradation in these cells (Supplementary Figure S7E-F). It appears that MS177 displayed higher anti-growth efficacies in AR-positive prostate cancer cells compared to those AR-negative ones (Figure 7A). MS177 also showed a superior effect on inhibiting proliferation of late-stage, highly plastic prostate cancer cells known to be resistant to antiandrogen therapy(36,37) (Supplementary Figure S7D), indicating a broader application of MS177 to various prostate cancer stages. Furthermore, MS177 was much more potent in inhibiting 22Rv1 cell growth than a larger panel of existing catalytic inhibitors of EZH2, including UNC1999(38,39), CPI-1205(40), EPZ-6438(41), and GSK126(42), and the EED degrader UNC6852 (Figure 7E-F and Supplementary Figure S7G). In addition, MS177 dose-dependently inhibited colony-forming capabilities (Figure 7G) and induced apoptosis, as assayed by immunoblotting for apoptosis markers (Figure 7H), in 22Rv1 cells, whereas the non-PROTAC controls, MS177N1 or C24, had little effect (Figure 7G-H). Together, the EZH2-targeting PROTAC MS177 robustly induces growth inhibition in a wide range of human prostate cancer cell lines and importantly, its cancer-killing effects are superior to those of EZH2 enzymatic inhibitors.

**Figure 7.**
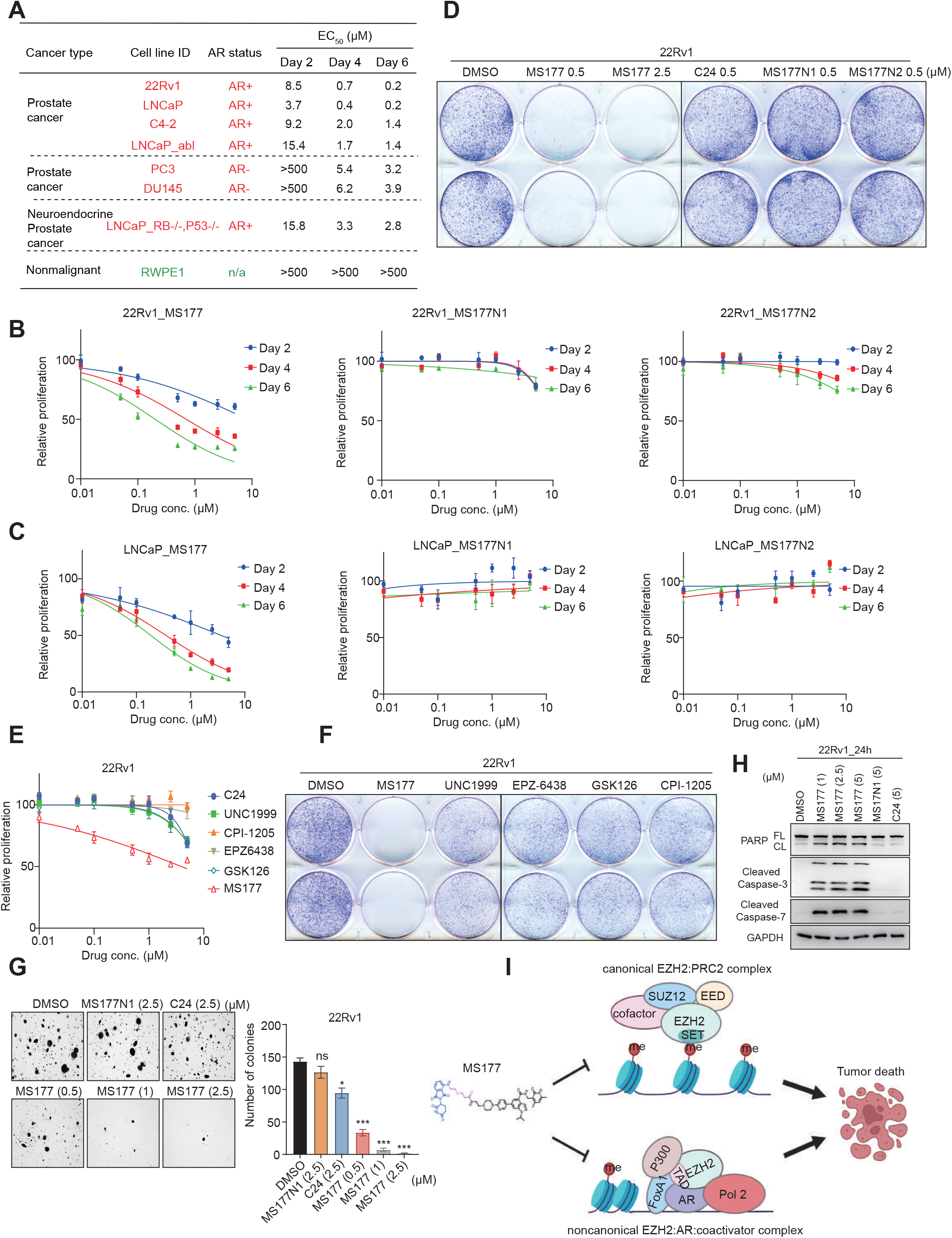
MS177 exhibits much more potent anti-proliferation effects in prostate cancer cell lines, compared to the enzymatic inhibitors of EZH2. **(A)** EC_50_ values of MS177 in the indicated prostate cell lines after a 2, 4 and 6-day treatment. EC50 values are the means of three independent experiments. **(B-C)** Plots showing the growth inhibitory effects of various used concentrations (x-axis; in the log10 converted values) of either MS177, MS177N1 or MS177N2 in 22Rv1 (**B**) and LNCaP (**C**) cells, treated for 2, 4 or 6 days. Y-axis shows relative growth after normalization to DMSO-treated cells (n= 3 independent treatment experiments; presented as the mean ± SD). **(D)** Colony formation assay using 22Rv1 cells treated with different doses of MS177 or 0.5 μM of C24, MS177N1 or MS177N2, compared to DMSO. **(E)** Plots showing the growth inhibitory effects of various used concentrations (x-axis; in the log10 converted values) of MS177 or various EZH2 inhibitors in 22Rv1 cells treated for 2 days. Y-axis shows relative cell growth after normalization to DMSO-treated cells (n= 3 independent treatment experiments; presented as the mean ±SD). **(F)** Colony formation assay using 22Rv1 cells treated with 2.5 μM of MS177 or the indicated EZH2 enzymatic inhibitors, compared to DMSO. **(G)** Representative views of soft agar-based assay (left panel) and quantifications of colony formation (right panel; colony numbers counted by ImageJ and presented in average ± SD of two independent experiments) using 22Rv1 cells, treated with DMSO, 2.5 μM of C24 or MS177N1, or three indicated concentrations of MS177 (n = 2 independent treatment experiments; presented as the mean ± SD). **(H)** Immunoblotting for the indicated apoptotic markers in 22Rv1 cells, treated with either 5 μM of MS177N1 or C24 or different indicated concentrations of MS177, compared to DMSO, for 24 hours. **(I)** A model showing that EZH2 forms canonical (EZH2:PRC2; top) and noncanonical (EZH2:AR:co-activators; bottom) complexes, both of which mediate tumorigenicity of advanced prostate cancer. The EZH2-targeting PROTAC MS177 simultaneously suppresses oncogenic circuits enforced by the above two of EZH2-associated onco-complexes, as well as those enforced by AR/AR-V7 signaling, thus leading to slowed proliferation and death of prostate tumor cells.

## Discussion

Increasing evidence pointed to multifaceted functions of EZH2 in cancer. In particular, EZH2 forms the so-called EZH2-solo sites in a PRC2-independent fashion and binds AR, eliciting the noncanonical gene-activation effects in prostate cancer(18). In this report, we unveiled, for the first time, (i) that EZH2 utilizes a hidden EZH2^TAD^ to form interactions with AR and/or AR’s constitutively-active variant AR-V7; here, we further mapped the EZH2^TAD^-interacting interfaces to at least two unstructured regions of AR— so-called polyQ- and polyG-containing sequences(43,44) with a property of intrinsically disorganized protein regions(45). These results suggest a multivalent protein-protein interaction involving un-structured protein sequences (TADs themselves often being unstructured) and possibly, liquid-liquid phase separation, which merits future studies. (ii) Integrated genomic profiling (CUT&RUN, ATAC-seq and RNA-seq), in combination with point mutagenesis of EZH2^TAD^, clearly demonstrated that the EZH2^TAD^-mediated association with AR/variants is crucial for establishment of the EZH2-solo:AR/AR-V7 cobinding pattern, for the transcriptional activation of their common targets, and for the potentiation of advanced CRPC cell growth in vitro and in xenografted mouse models. (iii) Furthermore, we show the higher expression of EZH2:AR:AR-V7 co-upregulated transcripts correlated with poorer outcomes of prostate cancer patients, thus demonstrating a clinical relevance for the oncogenic pathway studied by this work. (iv) Using a battery of biochemical, CUT&RUN and RNA-seq assays, we also observed that MS177, an EZH2-targeting PROTAC degrader, efficiently causes on-target degradation of EZH2, as well as EZH2-associated canonical (EZH2:PRC2) and noncanonical (EZH2:AR:AR-V7) onco-complexes in prostate tumor cells; here, we conducted careful MOA studies to show that such MS177-mediated degradation of AR/variant indeed relies on EZH2 binding by MS177 (as demonstrated by EZH2 depletion assay) and relies on the EZH2^TAD^-mediated AR interaction (demonstrated by using AR-interaction-defective mutant of EZH2^TAD^). (v) Lastly, MS177 is superior to existing catalytic inhibitors of EZH2 in suppressing the in vitro growth of prostate cancer cells. We have also attempted to assess in vivo efficacy of MS177 in a 22Rv1 cell xenografted mouse model; however, we could not achieve sufficient exposure levels for MS177 in this animal model (data not shown). Thus, we could not pursue this direction further with MS177. Optimization of MS177 into an improved EZH2 PROTAC degrader with better in vivo pharmacokinetic (PK) properties is needed to demonstrate in vivo efficacy for this therapeutic approach.

Interplays between EZH2 and AR/variants can be complex. Previous studies showed that EZH2 directly binds the AR gene promoter in a PRC2-independent manner to activate the transcription levels of AR and AR targets (such as PSA) and that EZH2 KD led to a significantly downregulated AR protein level(20,24,32). In this study, we carefully ruled out involvement of such a reported pathway in the MS177-induced degradation of AR/variants— indeed, within a short period-of-time of MS177 treatment (within 24 hours), we found the AR/variant mRNA levels unaltered and yet, a vast majority of AR/variant proteins were already subjected for ubiquitination and proteasomal degradation. Furthermore, given the existence of various cofactors for AR/variants, it is conceivable that only a subset of AR/variants in cells forms interaction with EZH2 at a given time, a notion also supported by their common but also distinctive genome-binding patterns. Thus, the observed dramatic and global effect by EZH2-targeting PROTAC (MS177) on AR/variant degradation is most likely due to the fastacting and re-cycling nature of MS177, which operates to essentially ‘drain’ the majority of cellular AR/variant proteins (with a half-life of approximately 6-7 hours(46)), in repeated cycles for degrading target/partners, from various AR/variant-containing complexes in prostate tumor. It appears that MS177 is more potent in the treatment of the AR-positive prostate tumor cells than those AR-negative ones, further pointing to the relevance of AR-degradation effects elicited by MS177. Please note that the EZH2^TAD^-mediated noncanonical gene-activation function may act in parallel with other EZH2-directed activities in prostate cancer— for example, EZH2 was recently shown to regulate 2’-O-Methylation of ribosomal RNA by directly interacting with fibrillarin, thereby promoting global protein translation of prostate cancer cells(21); moreover, EZH2 was also shown to bind to mutant p53 mRNAs, increasing their stability and capindependent protein translation in a methyltransferase-independent manner(47). Furthermore, EZH2 was reported to methylate FOXA1, leading to stabilization of this pioneer factor and activation of cell cycle-related genes(48). In theory, EZH2-targeting PROTACs represent a potentially attractive strategy for completely eliminating all multifaceted functions of EZH2 during prostate oncogenesis, which awaits further investigation.

## Data Availability

Genomic dataset of this study, including CUT&RUN and RNA-Seq, have been deposited in NCBI Gene Expression Omnibus (GEO) database under the accession number GSE205107. Human prostate cancer datasets were derived from TCGA Research Network (http://cancergenome.nih.gov/). Publicly available datasets used in the work were from NCBI GEO accession numbers GSE99378 (ATAC-seq data of 22Rv1 cells), GSM2827408 and GSE85558 (ChIP-seq for H3K27ac in 22Rv1 cells), GSE94013 (ChIP-seq for BRD4, AR and ARV7 in 22Rv1 cells, cultured under either ligand-stripped [mock] or ligand-stimulated [DHT] condition), GSE39459 (ChIP-seq for EZH2, H3K27me3, AR, H3K4me2, H3K4me3 and RNA Pol II in LNCaP-abl cells), GSE28950 (ChIP-seq for EZH2 and AR in VCaP cells), GSE14092 (ChIP-seq for H3K27me3 in VCaP cells), GSE123204 (ChIP-seq for EZH2 in PC3 cells), GSE57498 (ChIP-seq for H3K27me3 in PC3 cells), GSE135623 (ChIP-seq for EZH2 in DU145 cells), GSE82260 (ChIP-seq for H3K27me3 in DU145 cells), and GSE96652 (ChIP-seq for FOXA1 in 22Rv1 cells). Other data supporting the findings of this study are available upon request.

## Acknowledgments

We thank members of the Cai, Jin and Wang laboratories and X. Liu for technical supports and helpful discussion. We thank UNC core facilities, including Tissue Culture Facility, High-throughput Sequencing Facility (HTSF), Bioinformatics Core Facility and Animal Studies Core Facility, for professional assistance and support of this work.

