## supplemental materials for "A cryptic transactivation domain of EZH2 binds AR and AR’s splice variant promoting oncogene activation and tumorous transformation"

**Supplementary Table 1. RNA-seq identifies differentially expressed genes (DEGs) significantly downregulated in 22RV1 cells after EZH2 knockdown (KD; n= 2 biological replicate per group).**

**Supplementary Table 2. A list of the 130 genes significantly co-upregulated by full-length AR (AR-FL), AR-V7 and EZH2 in 22Rv1 CRPC cells, as determined by RNA-seq after KD of each gene (shAR-FL, or shAR-V7 or shEZH2) relative to mock (shEV; n= 2 biological replicate per group).**

**Supplementary Table 3. RNA-seq identifies DEGs showing significant up or down-regulation in 22RV1 cells after the treatment with 2.5 μM of MS177, C24 or MS177N1, relative to DMSO, for 24 hours (n= 2 biological replicate per group).**

**Supplementary Table 4. Sequence information for primers used in this study.**

### SUPPLEMENTARY FIGURES

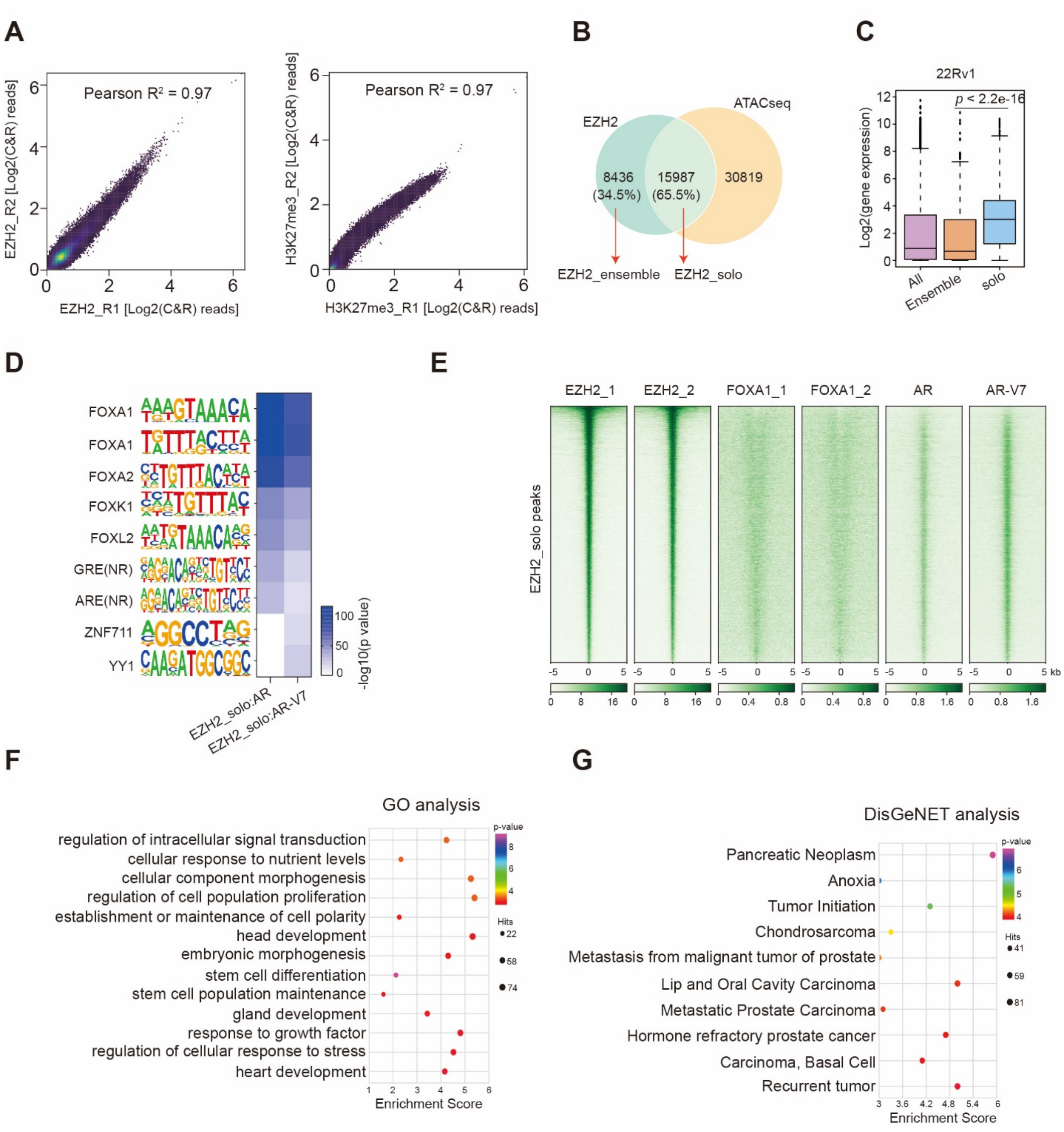


**Supplementary Figure S1. Besides its canonical H3K27me3-cobound EZH2:PRC2 sites, EZH2 also binds genomic sites that are characterized by the gene-activation-**

**associated chromatin markers and colocalization with AR or AR-V7 in prostate cancer.**

**(A)** Pearson correlation analysis of replicated EZH2 (left) or H3K27me3 (right) CUT&RUN signals in 22Rv1 cells (n=2 independent experiments).

**(B)** Venn diagram showing the indicated EZH2-solo or EZH2-ensemble peaks, determined by overlapping patterns of EZH2 CUT&RUN and ATAC-seq peaks.

**(C)** Box plot showing overall gene expression of all genes (left) and those associated with EZH2-ensemble (middle) or EZH2-solo (right) peaks in 22Rv1 cells. The boundaries of box plots indicate the 25th and 75th percentiles, the center line indicates the median, and the whiskers (dashed) indicate 1.5× the interquartile range. Paired two-sided t-test.

**(D)** Heatmap showing the enrichment of the indicated motifs at the EZH2-solo:AR (left column) or EZH2-solo:AR-V7 peaks (right column) in 22Rv1 cells.

**(E)** Heatmap for EZH2 (in duplicate), FOXA1 (in duplicate), AR and AR-V7 signals, ± 5 kb from the centers of EZH2-solo sites, in 22Rv1 cells.

**(F-G)** Gene ontology (GO) (**F**) and enrichment of the DisGeNet category (**G**) analyses using the targets commonly bound by EZH2-solo, AR and AR-V7 in 22Rv1 cells.

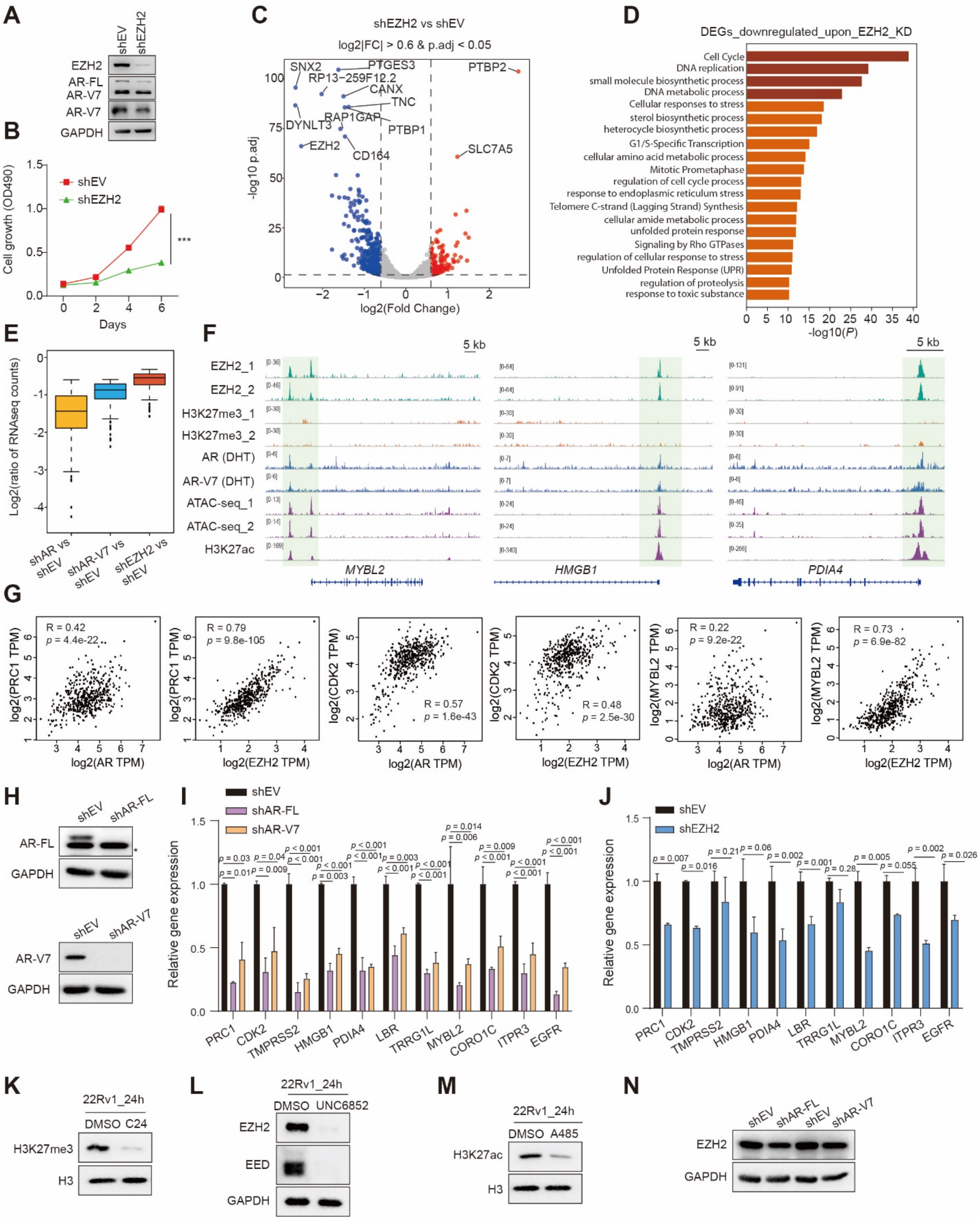

A
shEV
shEZH2
EZH2
AR-FL
AR-V7
AR-V7
GAPDH
B
shEV
shEZH2
Cell growth (OD490)
Days
***
C
shEZH2 vs shEV
log2|FC| > 0.6 & p.adj < 0.05
SNX2
PTGES3
RP13−259F12.2
CANX
TNC
DYNLT3
RAP1GAP
PTBP1
EZH2
CD164
PTBP2
SLC7A5
-log10 p.adj
log2(Fold Change)
D
DEGs_downregulated_upon_EZH2_KD
Cell Cycle
DNA replication
small molecule biosynthetic process
DNA metabolic process
Cellular responses to stress
sterol biosynthetic process
heterocycle biosynthetic process
G1/S-Specific Transcription
cellular amino acid metabolic process
Mitotic Prometaphase
regulation of cell cycle process
response to endoplasmic reticulum stress
Telomere C-strand (Lagging Strand) Synthesis
cellular amide metabolic process
unfolded protein response
Signaling by Rho GTPases
regulation of cellular response to stress
Unfolded Protein Response (UPR)
regulation of proteolysis
response to toxic substance
-log10(P)
E
Log2(ratio of RNAseq counts)
shAR vs shEV
shAR-V7 vs shEV
shEZH2 vs shEV
F
5 kb
EZH2_1
EZH2_2
H3K27me3_1
H3K27me3_2
AR (DHT)
AR-V7 (DHT)
ATAC-seq_1
ATAC-seq_2
H3K27ac
MYBL2
HMGB1
PDIA4
G
R = 0.42
p = 4.4e-22
log2(PRC1 TPM)
log2(AR TPM)
R = 0.79
p = 9.8e-105
log2(PRC1 TPM)
log2(EZH2 TPM)
R = 0.57
p = 1.6e-43
log2(CDK2 TPM)
log2(AR TPM)
R = 0.48
p = 2.5e-30
log2(CDK2 TPM)
log2(EZH2 TPM)
R = 0.22
p = 9.2e-22
log2(MYBL2 TPM)
log2(AR TPM)
R = 0.73
p = 6.9e-82
log2(MYBL2 TPM)
log2(EZH2 TPM)
H
shEV
shAR-FL
AR-FL
GAPDH
shEV
shAR-V7
AR-V7
GAPDH
I
shEV
shAR-FL
shAR-V7
Relative gene expression
PRC1
CDK2
TMPRSS2
HMGB1
PDIA4
LBR
TRRG1L
MYBL2
CORO1C
ITPR3
EGFR
J
shEV
shEZH2
Relative gene expression
PRC1
CDK2
TMPRSS2
HMGB1
PDIA4
LBR
TRRG1L
MYBL2
CORO1C
ITPR3
EGFR
K
22Rv1_24h
DMSO
C24
H3K27me3
H3
L
22Rv1_24h
DMSO
UNC6852
EZH2
EED
GAPDH
M
22Rv1_24h
DMSO
A485
H3K27ac
H3
N
shEV
shAR-FL
shEV
shAR-V7
EZH2
GAPDH

**Supplementary Figure S2. EZH2, AR and AR-V7 cooperate to activate transcription of a set of the clinically relevant oncogenes in prostate cancer.**

**(A-B)** Immunoblotting for the indicated protein **(a)** and in vitro growth of 22Rv1 cells **(b)** after EZH2 knockdown (KD; shEZH2). AR-FL, full-length AR.

**(C)** Volcano plot showing the DEGs, either up- (red) or down-regulated (blue) after EZH2 KD relative to mock in 22Rv1 cells. Top significant genes altered were highlighted.

**(D)** GO analysis of the DEGs downregulated after EZH2 KD in 22Rv1 cells.

**(E)** Box plots showing log2-converted ratios for the indicated RNA-seq sample comparisons by using the 130 genes identified in main Fig 2a to be coactivated by EZH2, AR and AR-V7. The boundaries of box plots indicate the 25th and 75th percentiles, the center line indicates the median, and the whiskers (dashed) indicate 1.5× the interquartile range.

**(F)** IGV views of the indicated factor at *MYBL2, HMGB1* and *PDIA4* in 22Rv1 cells.

**(G)** Spearman correlation plots for the mRNA expression levels of EZH2, AR and the indicated EZH2-solo:AR:AR-V7 targets in the TCGA prostate cancer cohort.

**(H)** Immunoblotting for AR and AR-V7 followed by AR-FL (top) or AR-V7-specific (bottom) KD in 22Rv1 cells. *, nonspecific bands.

**(I-J)** RT-qPCR for the indicated EZH2-solo:AR:AR-V7 co-targeted gene in 22Rv1 cells following specific KD of AR-FL or AR-V7 (**i**) or EZH2 (**j**). The y-axis shows averaged signals after normalization to those of GAPDH and to mock-treated (n = 3; mean ± s.d.; unpaired two-tailed Student's t-test).

**(K-M)** Immunoblotting of the indicated protein in 22Rv1 cells, treated with 2.5 µM of C24 (**K**), UNC6852 (**L**) or A-485 (**M**) relative to DMSO for 24 hours.

**(N)** EZH2 immunoblotting in 22Rv1 cells following by specific KD of AR-FL or AR-V7.

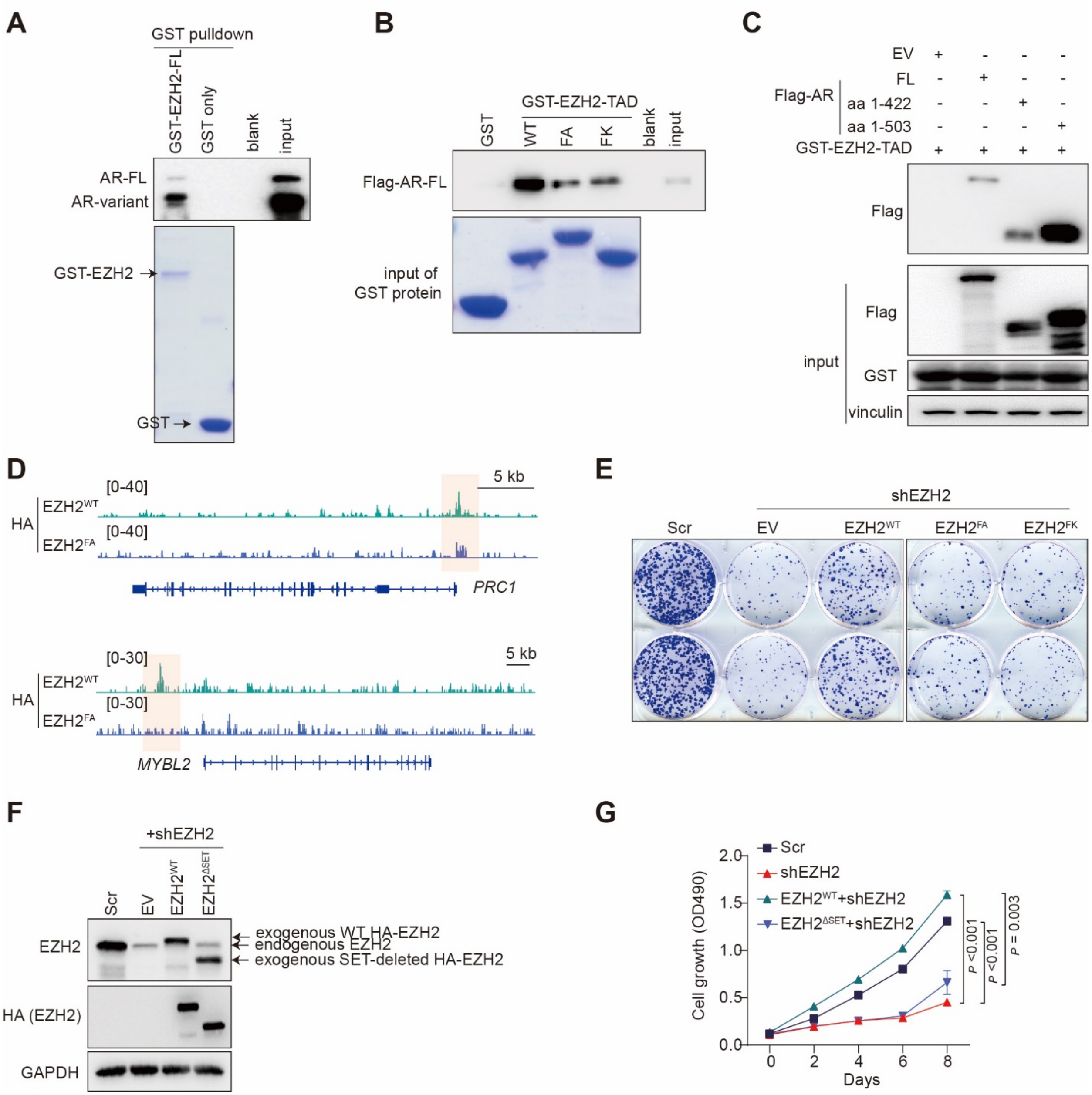


**Supplementary Figure S3. Interaction between EZH2$^{TAD}$ and AR is required for establishment of EZH2-solo binding at AR sites and for the malignant growth of prostate cancer cells.**

**(A)** GST pull-down assays using the recombinant GST protein, either GST alone or GST-EZH2 fusion, and the total lysate of 22Rv1 cells, followed by anti-AR immunoblotting (top). Bottom panel shows the GST protein input.

**(B)** GST pull-down using the indicated recombinant GST-fusion protein, either GST alone or that fused to wild-type (WT) or TAD-mutated (FA or FK) form of $EZH2^{TAD}$, and the total lysate of 293T cells transfected with the Flag-tagged full-length AR (AR-FL), followed by anti-Flag immunoblotting (top). Bottom panel shows the GST protein input.

**(C)** GST pull-down using the purified GST protein fused to WT $EZH2^{TAD}$ and the total lysate of 293T cells transfected with either empty vector (EV) or the indicated Flag-AR, followed by anti-Flag immunoblotting (top). Bottom panel shows the inputs.

**(D)** IGV views for the binding of HA-tagged EZH2, either WT or TAD-mutated (FA), at the indicated EZH2-solo:AR:AR-V7 targets (*PRC*1 and *MYBL2*) in 22Rv1 cells.

**(E)** Colony formation using the 22Rv1 cells pre-rescued with EV or the indicated exogenous HA-EZH2 (shEZH2-resistant; either WT or TAD-transactivation-dead mutant [FA or FK]), followed by depletion of endogenous EZH2 by shRNA (shEZH2) or mock treatment (scramble or Scr).

**(F-G)** Immunoblotting for the indicated exogenously-expressed HA-tagged EZH2 (**F**; either WT or a SET-deleted form; shEZH2-resistant) and measurement for in vitro growth (**G**) of the 22Rv1 cells pre-rescued with exogenous HA-EZH2, followed by mock treatment (Scr) or depletion of endogenous EZH2 (shEZH2). For **G**, n = 3, mean ± s.d.; unpaired two-tailed Student's t-test was used.

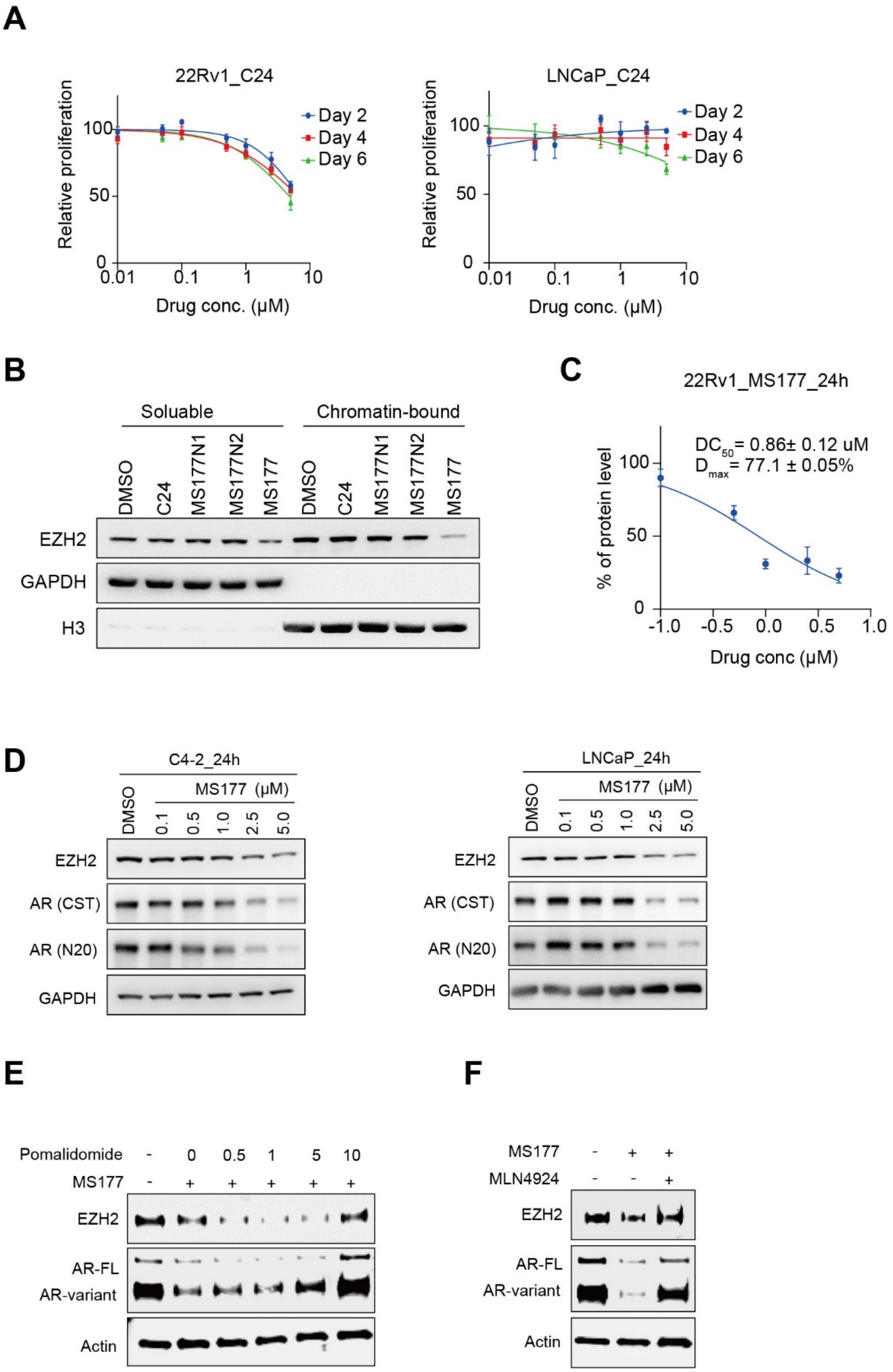
A
22Rv1_C24
LNCaP_C24
Relative proliferation
Day 2
Day 4
Day 6
Drug conc. (μM)
B
Soluable
Chromatin-bound
DMSO
C24
MS177N1
MS177N2
MS177
EZH2
GAPDH
H3
C
22Rv1_MS177_24h
DC50= 0.86± 0.12 uM
Dmax= 77.1 ± 0.05%
% of protein level
Drug conc (μM)
D
C4-2_24h
LNCaP_24h
MS177 (μM)
DMSO
0.1
0.5
1.0
2.5
5.0
EZH2
AR (CST)
AR (N20)
GAPDH
E
Pomalidomide
MS177
EZH2
AR-FL
AR-variant
Actin
F
MS177
MLN4924
EZH2
AR-FL
AR-variant
Actin

**Supplementary Figure S4. MS177, the EZH2-targeting PROTAC, mediates on-target degradation of EZH2, as well as EZH2-associated canonical (EZH2:PRC2) and non-canonical (EZH2:AR/AR-V7) complexes, in prostate cancer.**

**(A)** Plots showing growth inhibitory effects of various used concentrations of C24 (x-axis; in the log10 converted values) in 22Rv1 (left) and LnCaP (right) cells, treated for 2, 4 or 6 days. Y-axis shows relative cell growth after normalization to DMSO-treated cells (n= 3 independent treatment experiments; presented as the mean ±SD).

**(B)** Immunoblotting for EZH2, either nucleoplasmic (left) or chromatin-bound (right), in 22Rv1 cells after a 24-hour treatment with DMSO or 2.5 μM of C24, MS177N1, MS177N2 or MS177. GAPDH and histone H3 act as controls of cell fractionation.

**(C)** Measurement of $DC_{50}$ value of MS177 post-treatment of 22Rv1 cells, based on immunoblotting quantifications with ImageJ from two independent experiments. Mean ± s.d.

**(D)** Immunoblotting for EZH2 and AR (probed with two different antibodies, CST or N20) in C4-2 and LNCaP cells, treated with the indicated concentration of MS177, versus DMSO, for 24 hours.

**(E-F)** Immunoblotting for EZH2 and AR in 22Rv1 cells pre-treated with either DMSO or different concentrations of pomalidomide (**E**) or 0.4 μM of MLN4924 (**F**) for 2 hours, followed by an additional 24-hour treatment with 2.5 μM of MS177.

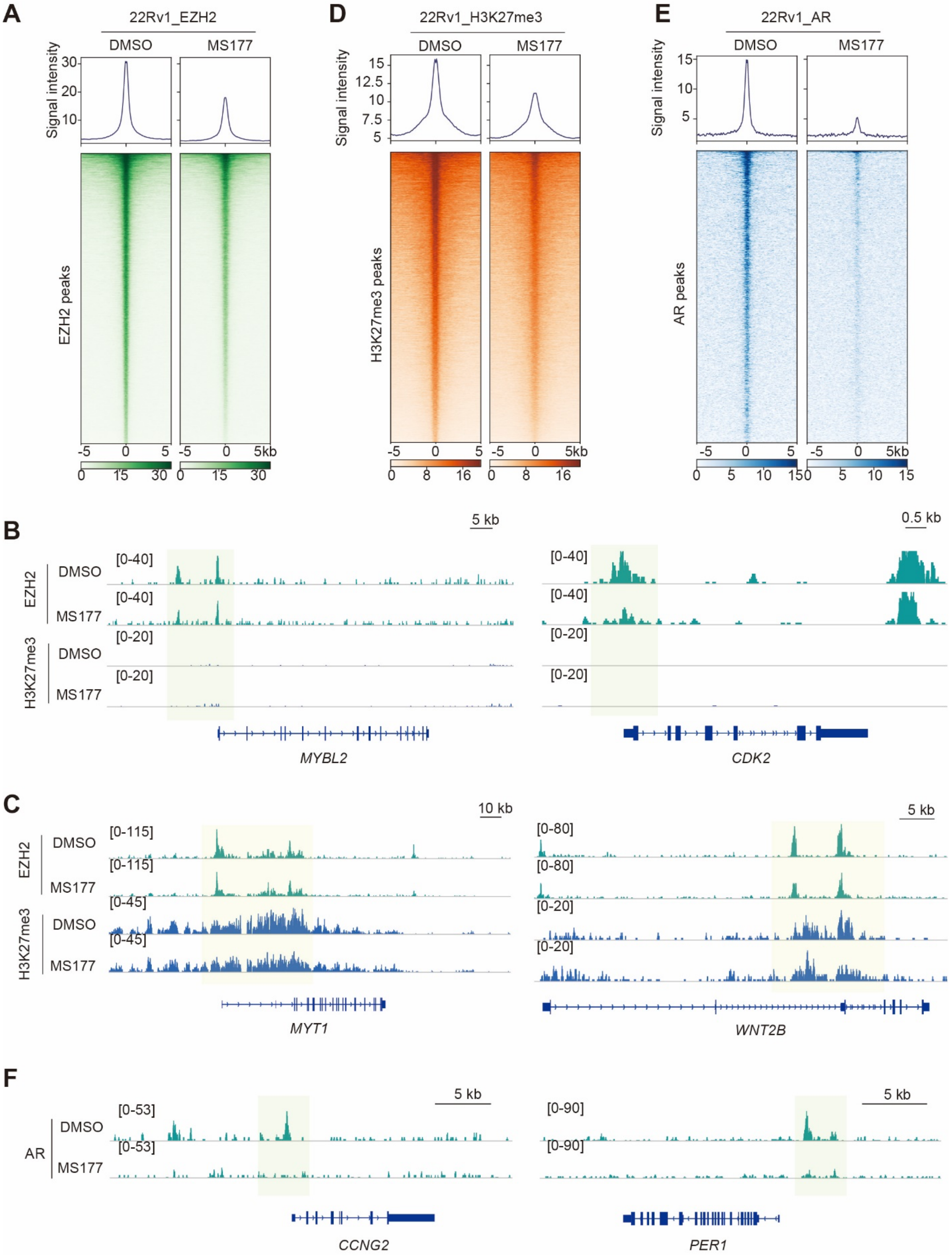

A
22Rv1_EZH2
DMSO
MS177
Signal intensity
EZH2 peaks
D
22Rv1_H3K27me3
H3K27me3 peaks
E
22Rv1_AR
AR peaks
B
EZH2
H3K27me3
MYBL2
CDK2
C
MYT1
WNT2B
F
AR
CCNG2
PER1

**Supplementary Figure S5. CUT&RUN-based profiling demonstrates effects by MS177 on decreasing genomic binding of both EZH2 and AR/AR-V7 in prostate cancer.**

**(A,D,E)** Average intensities (top panel) and heatmaps (bottom panel) for CUT&RUN signals (normalized against spike-in controls and sequencing depth) of EZH2, H3K27me3 and AR, ± 5 kb around the centers of EZH2 (**a**), H3K27me3 (**d**) and AR (**e**) peaks in 22Rv1 cells, treated with DMSO or 2.5 μM of MS177 for 24 hours.

**(B-C)** IGV views for EZH2 and H3K27me3 CUT&RUN signals (spike-in control and depth normalized) at the indicated EZH2-solo **(B;** *MYBL2* and *CDK2***)** or EZH2:PRC2-emsemble targets **(C;** *MYT1* and *WNT2B***)** in 22Rv1 cells, treated with DMSO or 2.5 μM of MS177.

**(F)** IGV views for CUT&RUN signals of AR binding (spike-in control and depth normalized) at *CCNG2* (left) and *PER1* (right) in 22Rv1 cells, treated with DMSO or 2.5 μM of MS177.

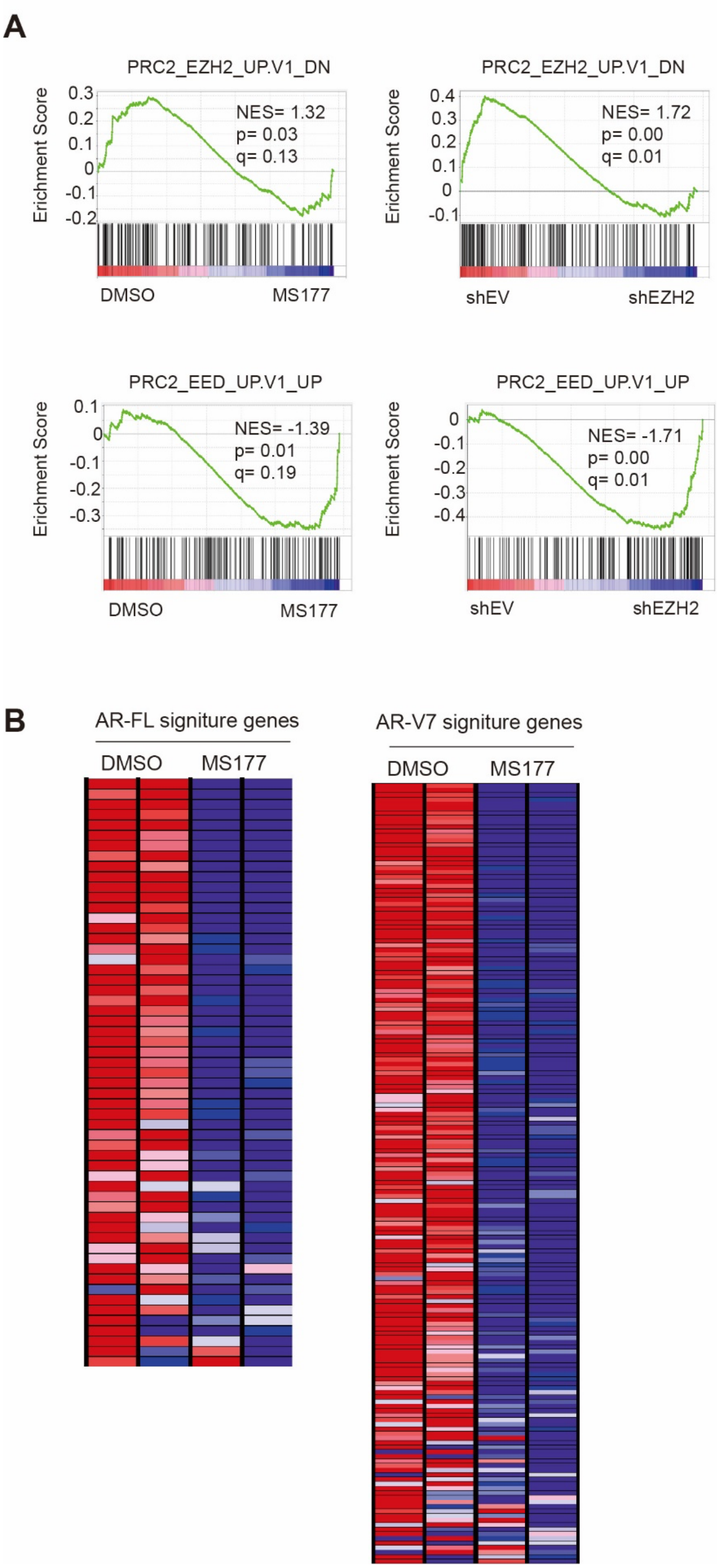
A
PRC2_EZH2_UP.V1_DN
Erichment Score
NES= 1.32
p= 0.03
q= 0.13
DMSO
MS177
PRC2_EZH2_UP.V1_DN
NES= 1.72
p= 0.00
q= 0.01
shEV
shEZH2
PRC2_EED_UP.V1_UP
NES= -1.39
p= 0.01
q= 0.19
DMSO
MS177
PRC2_EED_UP.V1_UP
NES= -1.71
p= 0.00
q= 0.01
shEV
shEZH2
B
AR-FL signiture genes
DMSO
MS177
AR-V7 signiture genes
DMSO
MS177

**Supplementary Figure S6. RNA-seq further substantiates unique and superior effects by MS177 on inhibiting both EZH2:PRC2- and AR/AR-V7-related oncogenic programs in prostate cancer.**

**(A)** GSEA revealing that MS177 treatment (left) and EZH2 KD (right) in 22Rv1 cells exhibited similar correlations with the increased expression of the PRC2-repressed genes. NES, normalized enrichment score.

**(B)** Heatmap showing the downregulation of AR-FL- (left) or AR-V7-activated (right) signature gene signatures in 22Rv1 cells following MS177 treatment relative to DMSO.

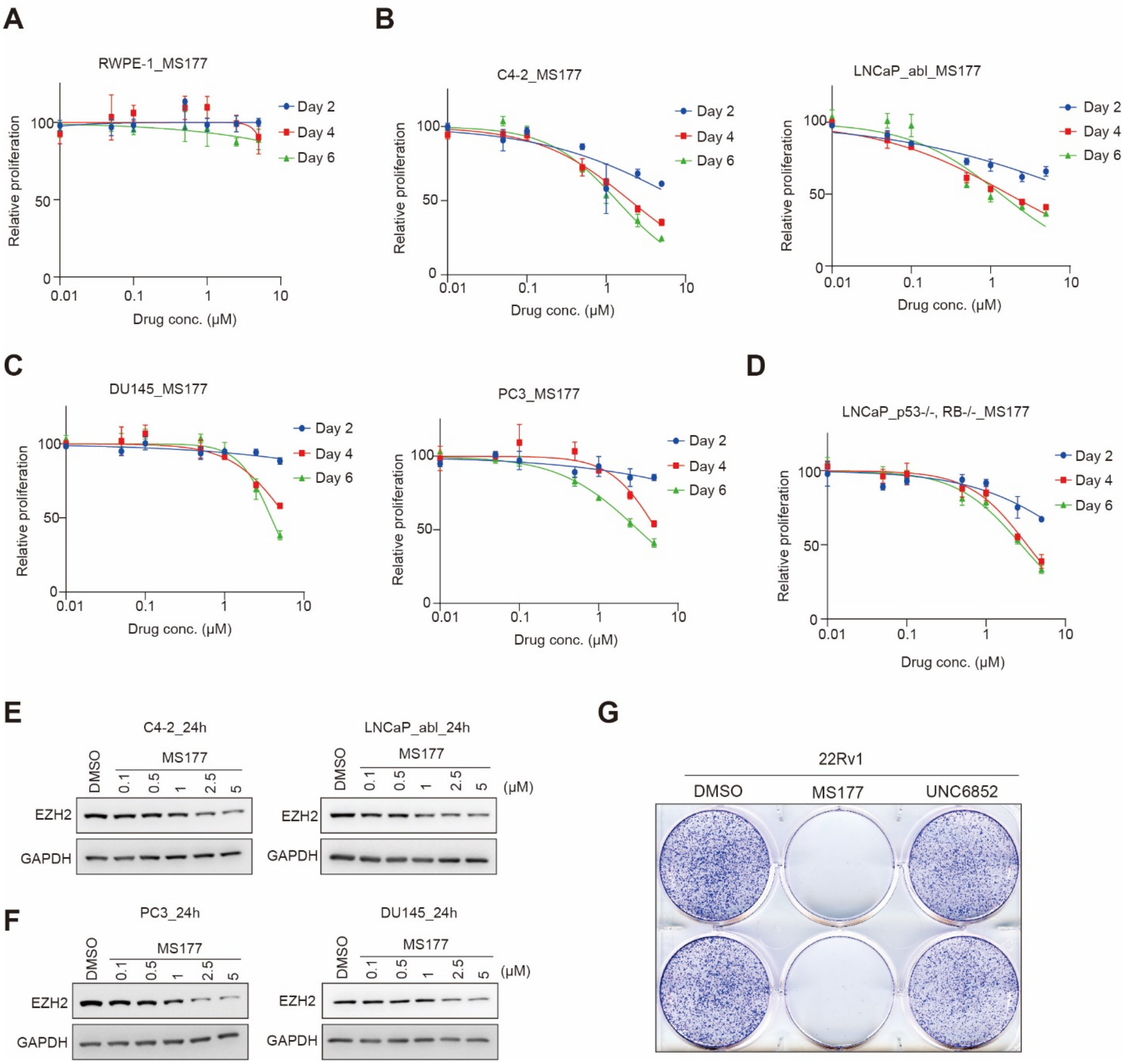


**Supplementary Figure S7. Compared to the EZH2 enzymatic inhibitors, MS177 elicits much more potent anti-tumor effects in prostate cancer cells.**

**(A-D)** Plots showing growth inhibitory effects of various used concentrations (x-axis; in the log10 converted values) of MS177 in RWPE1 cells **(A)**, AR-positive prostate cancer lines (namely, C42 and LNCAP_abl; **B**), those AR-negative ones (namely, DU145 and PC3; **C**), and LNCAP_P53-/-_RB-/- cells (**D**), treated for 2, 4 or 6 days. Y-axis shows relative cell growth after normalization to DMSO-treated cells (n= 3 independent treatment experiments; presented as the mean ±SD).

**(E-F)** Immunoblotting of EZH2 in different prostate cancer cell lines, either AR-positive (**E**) or AR-negative (**F**), treated with the indicated concentration of MS177 versus DMSO for 24 hours.

**(G)** Colony formation assay using 22Rv1 cells, treated with 2.5 μM of MS177 or UNC6852, compared to DMSO.
